# Moisture limits the diversity of nesting bees, wasps, and parasitoids in lying and standing deadwood

**DOI:** 10.64898/2026.04.10.717477

**Authors:** Massimo Martini, Matteo Dadda, Felix Fornoff, Heike Feldhaar, Arong Luo, Finn Rehling, Joshua E. Spitz, Michael Staab, Simon Thorn, Chao-Dong Zhu, Alexandra-Maria Klein

**Affiliations:** Chair of Nature Conservation and Landscape Ecology, Faculty of Environment and Natural Resources, University of Freiburg; Freiburg, Germany; State Key Laboratory of Zoological Systematics and Evolution, Institute of Zoology, Chinese Academy of Sciences; Beijing, China; Animal Population Ecology, Faculty of Biology, Chemistry & Earth Sciences, University of Bayreuth; Bayreuth, Germany; Animal Ecology and Trophic Interactions, Institute of Ecology, Leuphana University of Lüneburg; Lüneburg, Germany; Department of Biology, University of Marburg, Marburg, Germany; Hessian Agency for Nature Conservation, Environment and Geology, Biodiversity Centre, Gießen, Germany; Czech Academy of Sciences, Biology Centre, Institute of Entomology, České Budějovice, Czech Republic

**Keywords:** Ant exclusion experiments, deadwood-nesting Hymenoptera, biodiversity-ecosystem functioning, host-parasitoid interactions, standing and lying deadwood, substrate moisture, tree diversity experiments

## Abstract

Saproxylic insect community assembly is structured by deadwood and forest habitat gradients, as well as biotic interactions such as competition, predation, and parasitism. However, co-variation between abiotic and biotic conditions limits our ability to disentangle their contributions. Furthermore, a focus on beetles in temperate and boreal forests has left important taxonomic and geographic knowledge gaps. Here, we tested how experimentally manipulated tree diversity, deadwood position (lying vs. standing), and biotic interactions with a dominant antagonist (ant exclusion) structure communities of deadwood-cavity-nesting bees, wasps, and their parasitoids in a subtropical forest. Lying deadwood supported less diverse and abundant communities than standing deadwood while retaining approximately twice as much moisture. Moreover, host emergence declined along moisture gradients within each deadwood type. Together, these patterns identify substrate moisture as an important factor limiting bee and wasp communities nesting in deadwood – reflecting reduced brood production, survival, or both – and as a candidate driver for their positive association with standing deadwood in forests. By contrast, neither ant exclusion nor observed ant occurrence was associated with host or parasitoid responses. Because standing deadwood is typically scarce in managed forests, our findings support retaining or creating standing deadwood structures alongside lying deadwood and, more broadly, promoting dry and elevated nesting substrates for cavity-nesting bees and wasps in conservation and restoration measures.

**Graphical abstract:** 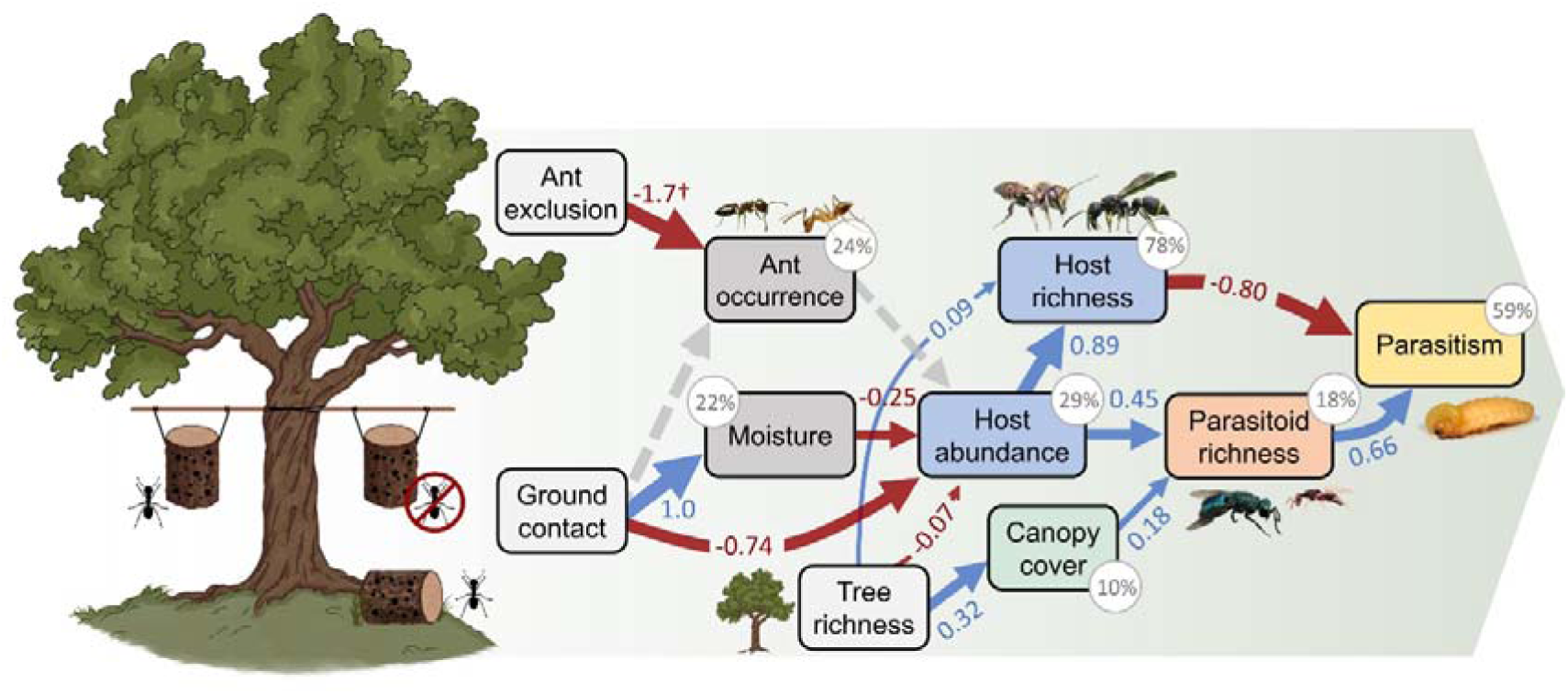

## Introduction

Saproxylic communities are structured by environmental constraints operating across hierarchical habitat scales (Neff et al., 2022). At the forest scale, tree diversity and composition, canopy cover, and total deadwood volume filter species from a regional pool (Seibold et al., 2015). Individual deadwood objects represent discrete habitat patches that are then colonized according to properties such as tree species identity, diameter, decay stage, and spatial positioning (Seibold et al., 2015). Species’ occurrence and relative abundance may be further shaped by biotic interactions including competition, facilitation, and trophic interactions among co-occurring taxa (Zou et al., 2023; Saine et al., 2025). In natural forests, these processes are difficult to disentangle because micro- and macrohabitat conditions, and biotic interactions often co-vary. Manipulative experiments that simultaneously capture environmental and biotic constraints across habitat scales remain scarce. Moreover, most work has focused on beetles and fungi in temperate or boreal forests, leaving important taxonomic and geographic gaps that limit generalization (Seibold et al., 2015).

Deadwood cavity-nesting bees and wasps (Hymenoptera: Apocrita) represent an ideal system to investigate mechanisms for saproxylic community assembly. These insects use deadwood as nesting substrate and occupy multiple trophic levels (pollinators, predators, and their parasitoids). As such, cavity-nesting Hymenoptera are highly sensitive to micro- and macrohabitat conditions because environmental filtering acts directly on their own requirements and indirectly through the availability of cavities, food, or hosts (Tscharntke et al., 1998). About 20% of solitary bees and wasp species nest in cavities, and up to one quarter of these are strongly associated with deadwood cavities, such as those created by wood-boring beetles (Ulyshen, 2018). Despite their diversity and ecological importance, these insects remain poorly studied (Ulyshen, 2018). Evidence from standardized trap nests (Fig. A1) indicates sensitivity to forest structural properties, including stand structural complexity and availability of standing deadwood (Rappa et al., 2023, 2024). Tree diversity, composition, and canopy cover shape communities based on forest specialization (Fornoff et al., 2021; Rappa et al., 2023; Wang et al., 2025). Moreover, local microclimatic and nesting conditions may strongly influence colonization and brood production (Hranitz et al., 2009; Mayr et al., 2020; Polidori et al., 2024), even outweighing surrounding landscape characteristics (Everaars et al., 2011). However, the mechanisms through which micro- and macrohabitat conditions jointly shape communities nesting in deadwood remain poorly understood.

The spatial positioning of deadwood represents a key microhabitat factor that can modify both abiotic conditions and biotic interactions (Law et al., 2019; Barrera-Bello et al., 2024). Deadwood lying on the forest floor typically retains more moisture, receives less solar exposure, may host different or more diverse fungal communities, and decomposes more rapidly than standing deadwood (Harmon et al., 2011; Oechler et al., 2025). These differences may affect nesting suitability and larval development in nesting Hymenoptera, which are sensitive to humidity gradients and associated fungal activity (Stephen, 1965; Flores et al., 1996). Indeed, bees and wasps colonize standing deadwood more frequently than lying deadwood and are generally more abundant and diverse in forests with greater availability of standing – but not lying – deadwood (Westerfelt et al., 2015; Rappa et al., 2023). However, direct experimental evidence of how deadwood position affects cavity-nesting Hymenoptera remains scarce, and the driving factors are not well described. Deadwood position may also influence community assembly indirectly by altering biotic interactions. Deadwood in contact with the ground may be more likely to be colonized by ants (Boucher et al., 2015), which commonly nest and forage in deadwood and can represent one of the most abundant animal groups in this substrate (Ulyshen, 2018). Ants can act as dominant antagonists in saproxylic systems, altering community structure and associated ecosystem functions (Law et al., 2019; Wu et al., 2021). In subtropical environments, ants frequently occupy trap nests or prey on bee and wasp brood, reducing nesting activity and offspring survival (Miyano & Yamaguchi, 2001). Conversely, parasites, parasitoids, and kleptoparasites attacking bee and wasp nests may benefit from more exposed and accessible nesting substrates such as standing deadwood (Hedgren, 2007). Both natural enemies and deadwood moisture may further be directly influenced by forest structure (Forrester et al., 2012; Staab & Schuldt, 2020). Variation in deadwood position and forest structure may therefore influence cavity-nesting Hymenoptera communities through combined effects on nesting substrate conditions and biotic interactions.

Here, within a large forest biodiversity experiment, we tested how experimentally manipulated microhabitat (deadwood position), macrohabitat (tree diversity), and biotic interactions with ants (ant exclusion) drive the diversity, composition, and parasitism of deadwood cavity-nesting Hymenoptera. We hypothesized that (1) deadwood position, and to a lesser extent tree diversity, will constrain the composition of cavity-nesting Hymenoptera communities; (2) hosts and parasitoid diversity and parasitism will be higher in standing compared to lying deadwood due to lower moisture and reduced ant disruption providing more optimal nesting conditions; and (3) host and parasitoid diversity will be positively associated with tree diversity and canopy cover, but macrohabitat effects are weaker than microhabitat. Our study addresses key taxonomic and geographic gaps and evaluates candidate factors associated with the assembly of non-obligate multitrophic communities in deadwood.

## Materials and methods

### Study sites and plots

This study was conducted as part of the MultiTroph project in the BEF-China experiment, located in subtropical southeast China in Jiangxi province (117°54’ E, 29°07’ N; Klein et al., 2026). It includes two replicated field sites (A and B, Fig. A2) situated 4 km apart from each other, approximately 20 ha each, and is currently the largest forest tree diversity experiment worldwide (Liu et al., 2026). Across the experiment, 566 study plots were established by planting seedlings from a pool of 40 native tree species in 2009 (site A) and 2010 (site B). Each plot measures 25.8 × 25.8 m and contains 400 trees planted in a 20 × 20 grid, with 1.29 m spacing. At each site, plots were randomly assigned a tree diversity of 1, 2, 4, 8, 16 or 24 planted species. Herbaceous vegetation and non-planted woody species are removed from all plots twice annually. The experimental tree species are known to directly and indirectly provide resources for cavity-nesting Hymenoptera, including pollen for bees (Zhang et al., 2026) and arthropod prey for wasps (Fornoff et al., 2019; Wang et al., 2019; Chen et al., 2023). See Bruelheide et al. (2014) for more details on the BEF-China experiment.

### Sampling design

We used standardized deadwood traps (Fig. A1) to sample cavity-nesting Hymenoptera, including bees, wasps, and their parasitoids. Each trap consisted of a cross-section of *Schima superba* Gardn. & Champ (Theaceae), a common local tree species present in the BEF-China experiment and producing intermediate-density wood. Deadwood was sourced from non-experimental trees outside the study plots in autumn 2023; all logs were dried together under ambient conditions over winter before being randomly assigned to plots and treatments in spring 2024. Logs measured ∼ 30 × 10 cm (length × diameter) and contained 44 cavities drilled horizontally into the side through the bark. Cavity diameters matched the known distribution used by local cavity-nesting Hymenoptera species (3-12 mm, see Table A1 for details; Staab et al., 2016; Fornoff et al., 2021). At each site, we selected 32 plots spanning six planted tree-species-richness levels: 16 monocultures, eight 2-species mixtures, four 4-species mixtures, two 8-species mixtures, one 16-species mixture, and one 24-species mixture (64 plots in total; Fig. A2).

In each plot, three deadwood traps were installed at a focal tree ≥ 9 m away from the plot edge. One trap was placed at the base of the tree to mimic lying deadwood. Two more traps were suspended at 1.3 m height from a horizontal metal bar strapped to the trunk to represent elevated sections of standing deadwood. Both standing and lying deadwood cavity orientation was horizontal to the gravitational plane. In one of the two standing traps, ants were excluded by applying resin-based insect glue on the supporting bar. Focal trees were chosen for adequate size to safely support the suspended deadwood. This design resulted in 192 deadwood traps in total (3 treatments × 32 plots × 2 sites). Each drilled log was paired with an adjacent undrilled log of the same type and size used to measure gravimetric moisture content (GMC, Fig. A1).

Deadwood traps were exposed in the field for 11 months (April 2024 - March 2025). Sticky resin barriers were inspected opportunistically throughout this time period and re-applied whenever visibly compromised. Traps were then carefully removed, enclosed in fine-mesh nylon bags, and suspended vertically outdoors under a rain shelter to collect emerging insects. Emergence vials containing 99% ethanol were attached to the bottom of each trap (Fig. A3) and monitored weekly for five months, until no further emergence was observed for 14 days. We identified all emerged bees and wasps to species or morphospecies based on specimens from the long-term BEF-China reference collection (Martini et al., 2026), available taxonomic keys, and consultation with expert taxonomists. Morphospecies (representing ∼7% of emerged insects) were used where the available taxonomic resources did not permit reliable assignment of formal species names. Voucher specimens are kept at the Institute of Zoology, Chinese Academy of Sciences (Beijing). Insects exploiting host nests were grouped as “parasitoids” based on life-history information. All nesting bees and wasps per deadwood object were treated as one “host” community, and all their parasitoids as a second community to test general responses of species richness and abundance (Staab et al., 2016). Based on the known life histories of the recorded parasitoid taxa, none were gregarious or killed multiple host individuals. Assuming that each parasitoid individual killed one host individual (or brood cell), we calculated apparent parasitism rate for each deadwood object as *P/(H+P),* where *P* is the number of emerged parasitoid individuals and *H* is the number of emerged hosts.

### Habitat variables

We measured several micro- and macrohabitat variables to assess their effect on insect community assembly. Macrohabitat (plot-level) characteristics included tree species richness, canopy cover, and coarse woody debris (CWD) volume. Canopy cover, which influences microclimate and light availability, was measured in August 2024. We took hemispherical pictures at 1.3m above ground with an 8mm fisheye lens perpendicular to the gravitational plane and calculated the percentage of sky area covered by plants using ‘Gap Light Analyzer’ v2.0 (Frazer et al., 1999). CWD volume was calculated by measuring the length and circumference of all deadwood pieces of diameter > 7 cm, including naturally-occurring deadwood objects and stumps remaining from previous commercial tree plantations (Dadda et al., 2026).

Microhabitat conditions included deadwood position, substrate moisture, and ant occurrence. To calculate wood moisture we cut a disk of 5 cm thickness from a random end of each non-drilled log (3/plot) immediately after removal from the field and measured its wet mass. Disks were oven-dried at 60 °C for 120 h and re-weighed to determine dry mass. Gravimetric moisture content was then calculated as the weight of the water in each deadwood disk relative to its dry mass. We used this measure as a proxy of moisture conditions of the adjacent deadwood trap.

Ants were quantified as biotic interactors at both micro- and macrohabitat scale. Plot-level ant activity was measured using Winkler litter extraction (Agosti et al., 2000). Four 1 m^2^ litter samples, collected at the corners of the central 10x10-tree area, were extracted into ethanol for 48 h. Ants were identified to species or morphospecies using taxonomic literature and reference material (following Staab et al., 2014). We calculated a plot-level activity index by summing species incidences across the four samples (maximum four occurrences per species). At the deadwood level, ant occurrence was recorded from the emergence samples and used as the primary predictor of host and parasitoid responses, indicating whether ants had accessed or occupied the nesting substrate. Ants were detected in three ant-exclusion traps (4.7% protocol failure rate), as fallen or growing branches likely created new access routes that bypassed the resin barriers (although entry pathways were not observed directly). These deadwood traps were included in (not excluded from) the analyses. We considered potential effects of plot geomorphology on deadwood moisture and ant occurrence by assigning each plot to one of three categories: (i) spur, ridge, or summit (23 plots); (ii) slope (29 plots); and (iii) hollow or valley (12 plots).

A small proportion of missing data was imputed prior to analysis using model-based predictions and residual variance for deadwood moisture (19 traps; 9.9% of data), CWD (4 plots; 6.25% data), and plot-level ant activity (3 plots; 4.7% of data; Methods B1). Imputed variables were only used as predictors to avoid loss of observations.

### Statistical analyses

We conducted all analyses using R v.4.6.1 (R Core Team, 2026).

### Sampling completeness and robustness of richness estimates

Before testing our hypotheses, we evaluated whether observed species richness adequately represented the sampled communities. For hosts and parasitoids separately, we constructed sample-based species-accumulation curves by randomly permuting the order of occupied deadwood traps 1000 times and calculated jackknife estimates of total richness (Oksanen et al., 2025). We then calculated abundance-based sample coverage for hosts at the overall, plot, and trap levels (Chao et al., 2020). Host coverage was consistently high (overall coverage = 0.99; mean plot coverage = 0.94; mean trap coverage = 0.95). As a sensitivity analysis, we repeated the main inferential host richness analysis using richness estimates standardized to a common trap coverage of 0.90. Resulting patterns were qualitatively consistent with those based on observed richness. We therefore retained observed host species richness as the primary response.

### Primary inference using generalized linear mixed models

Generalized linear mixed models (GLMMs) provided the primary test of our ecological hypotheses. Models were fitted using the ‘glmmTMB’ package (Brooks et al., 2025). We used response-specific models to test how deadwood treatment and forest environment affected host and parasitoid communities. Predictors measured at deadwood-level were treatment, moisture content, and ant occurrence, whereas plot-level predictors were tree species richness, canopy cover, and CWD volume. Many traps remained without nests, so host abundance was analyzed using a zero-inflated negative-binomial model that distinguished non-colonization from variation in abundance among colonized traps (Potts & Elith, 2006). Treatment and site were included in the zero-inflation component because they affected the probability of non-colonization (Fig. A4). Parasitoid abundance, richness, and apparent parasitism were analyzed only for traps with evidence of host brood, defined as successful host or parasitoid emergence (*n* = 127), thus removing structural zeroes.

To distinguish direct associations with richness from those arising through variation in abundance, richness models were fitted both with and without abundance from the corresponding trophic level. Parasitoid models also included host abundance to evaluate bottom-up associations, while the apparent-parasitism model included host and parasitoid richness to test for multitrophic diversity effects. In addition to these full models that tested marginal predictor effects, we fitted reduced models containing only the experimentally-manipulated deadwood and tree-richness treatments. These reduced models estimated their total effects without conditioning on potentially downstream variables such as moisture, abundance, or richness.

We also tested whether deadwood moisture was associated with treatment, tree richness, canopy cover, and plot geomorphology, and whether deadwood ant occurrence was associated with treatment, plot-level ant activity, and the same plot variables. Abundance, richness, occurrence, moisture, and apparent parasitism responses were fitted with negative binomial, Poisson (or generalized-Poisson when under-dispersed), binomial, beta, and beta-binomial distributions, respectively. Models included random intercepts for plots nested within sites. Random intercepts were omitted if variance approached zero and prevented model convergence (Table C1). Tree richness was log2-transformed and CWD was square-root-transformed. Host and parasitoid abundance were log(y +1)-transformed when used as predictors, and all continuous predictors were standardized to a mean of zero and standard deviation of one. When included as a predictor, moisture was centered and standardized separately within each deadwood treatment. Its coefficient therefore captured the association between the moisture gradient within each treatment and the response, rather than systematic moisture differences among treatments. For host abundance, we additionally fitted an alternative model using moisture standardized across the full dataset to assess the association with moisture across treatments. Model assumptions were evaluated using residual diagnostics, and multicollinearity was assessed using pairwise correlations (Fig. A5) and variance inflation (VIF < 3). Model sensitivity was assessed by refitting host occurrence and abundance with plot-level ant activity or log-transformed deadwood-level ant abundance instead of ant occurrence. We also repeated all community models using complete cases (*n* = 173), excluding imputed moisture and CWD values. Sensitivity analyses produced qualitatively-consistent results (Table D1-D3).

### Treatment and environmental effects on host community composition

We used multivariate analyses to determine whether deadwood treatments and forest conditions affected host community composition in addition to abundance and species richness. We tested variation in host community composition at the deadwood trap and plot levels using permutational multivariate analysis of variance (PERMANOVA) based on Bray–Curtis dissimilarities of square-root-transformed species abundances. At the trap level, we tested the effects of deadwood treatment, within-treatment moisture, and ant occurrence, restricting permutations within plots. At the plot level, we tested the effects of tree richness, canopy cover, and plot-level ant activity. We additionally repeated the plot-level analysis using binary (presence-absence) data to determine whether compositional patterns reflected species occurrence rather than differences in relative abundance. All analyses used 1,000 permutations (Oksanen et al., 2025). Prior to inference, we tested for differences in multivariate dispersion using parametric ANOVA and permutation-based PERMDISP tests (Table C2).

We visualized compositional patterns using distance-based redundancy analysis (dbRDA), constrained by deadwood treatment and tree richness, and extracted species scores to identify the taxa most strongly associated with these gradients. We complemented the ordination with indicator-species analyses for deadwood treatment and position (De Cáceres et al., 2010). Finally, we partitioned pooled presence-absence Sørensen dissimilarity between lying and standing deadwood into turnover and nestedness components (Baselga et al., 2025) to characterize the compositional difference. We did not analyze parasitoid community composition because data was too sparse to support reliable multivariate inference.

### Direct and indirect relationships using path analysis

We conducted path analyses using directed tests of separation with ‘piecewiseSEM’ to determine conditional associations between variables in the multi-variate framework (Shipley, 2009; Lefcheck, 2016). This analysis is complementary to the primary mixed models and allowed us to investigate support for a network of hypothesized direct and indirect pathways linking deadwood treatments and tree richness to Hymenoptera communities via altered nesting and forest conditions. We first constructed a maximal model with all hypothesized pathways, including potential effects of treatment and tree richness on the two trophic groups, bottom-up relationships from hosts to parasitoids, and associations of host and parasitoid diversity with realized apparent parasitism. Canopy cover, across-treatment deadwood moisture, and ant occurrence were included as potential intermediate pathways. We also specified non-directional correlations between variables where we assumed no causal ecological relationship, including ant occurrence with moisture content (both correlated with plot geomorphology), and apparent parasitism with host abundance (reflecting their mathematical dependence). To remain consistent with our main GLMMs, we fitted direct pathways using the same data subsets (e.g. excluding structural zeroes in parasitoid and parasitism component models). The direct path from tree richness to canopy cover was fitted using a plot-level dataset (*n* = 64). We assessed global model fit using Fisher’s C statistic derived from tests of directed separation, for which *p* > 0.05 indicates that the hypothesized network structure is not rejected (see maximal model configuration and results in Table C3). We additionally derived a parsimonious representation by sequentially removing unsupported pathways while monitoring AIC and global fit. For both the maximal and parsimonious models, we derived standardized coefficients for continuous pathways from matched Gaussian-standardization models with identical pathway structure (Methods B2). Pathway support and *p-*values were inferred from the original non-standardized models.

## Results

Overall, 26 cavity-nesting bee and wasp species (1099 individuals) and 13 parasitoid species (129 individuals) emerged from the deadwood traps (Table A2), representing 76% and 68% of the predicted richness, respectively (jackknife, Fig. A6). Standing deadwood supported 25 host and 11 parasitoid species (96% and 85% of total) across 128 traps, whereas lying deadwood supported 8 host species and two parasitoid species only (31% and 15%) across 64 traps. Overall, hosts emerged from 79% of standing compared to only 38% of lying traps, whereas parasitoids emerged from 38% and 8% of standing and lying traps, respectively.

Deadwood position strongly determined host and parasitoid diversity, whereas ant exclusion had no effect. Host and parasitoid abundance and species richness were consistently higher in standing compared to lying deadwood, but did not differ between standing deadwood with and without the ant exclusion protocol (Fig. 1A-D). Parasitism did not differ between deadwood treatments (Fig. 1E). Neither trophic level nor parasitism showed any association with observed ant occurrence on deadwood. (Table C1).

**Figure 1.**
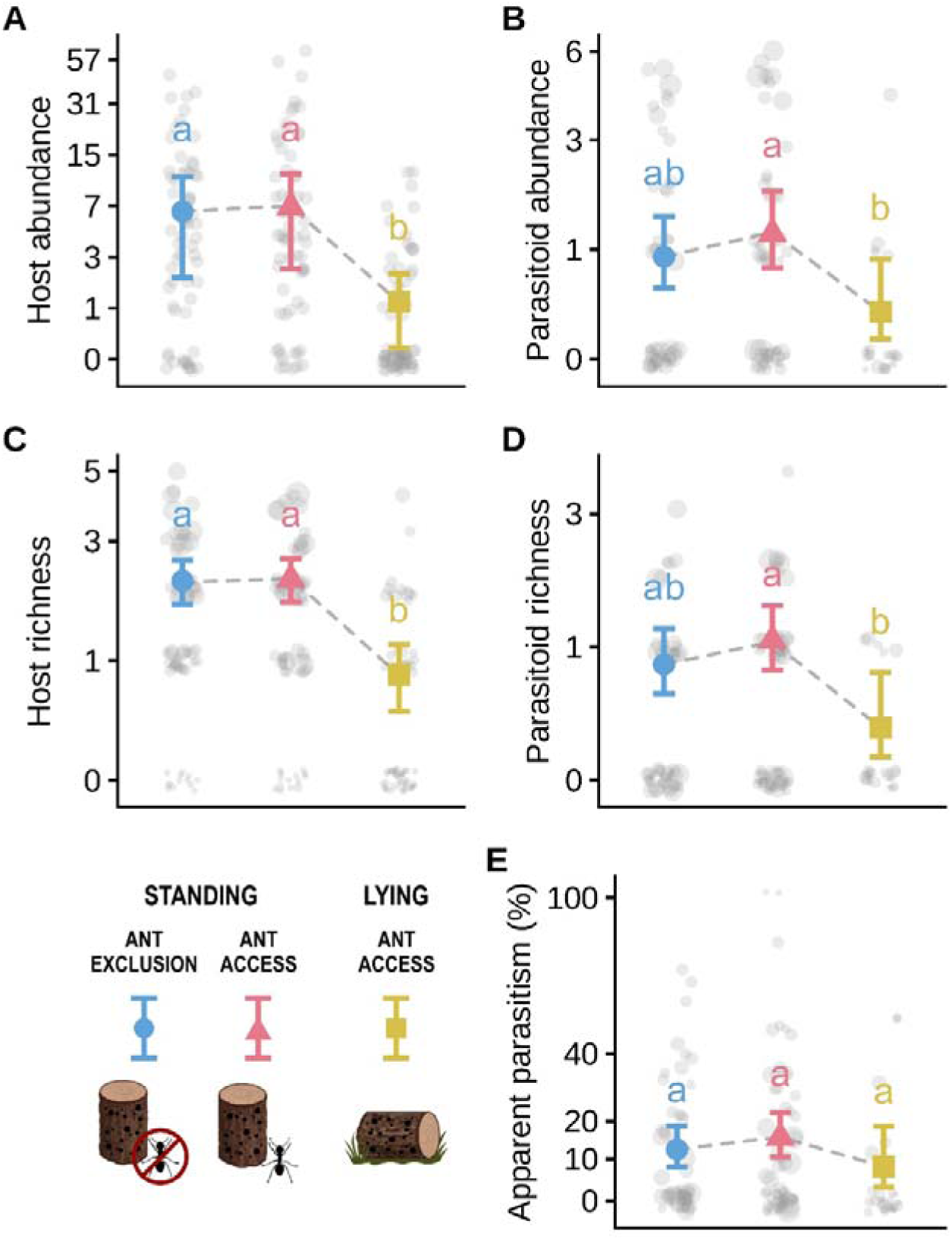
Saproxylic host (bees and wasps) and parasitoid abundance (A, B), species richness (C, D), and apparent parasitism rate (E) in standing deadwood with ant exclusion (blue circle), standing deadwood accessible to ants (red triangle), and deadwood lying on the ground (yellow square). Larger points in panels B-E indicate higher host abundance, symbols represent marginal effects (±95% CI) from GLMMs. Y-axes are displayed on a log(y + 1) scale for improved visualization, with tick labels shown in the original response units. Letters indicate significant differences based on Tukey-adjusted pairwise comparisons.

Deadwood position generated strong differences in substrate moisture: lying deadwood retained approximately twice as much moisture compared to standing deadwood (Fig. 2A). Path analyses were consistent with an indirect pathway linking lying deadwood to lower host and parasitoid diversity through higher substrate moisture (Fig. 3). Higher moisture from ground contact was associated with decreased host emergence (Fig. 4A) and consequently lower host and parasitoid richness. This negative association between substrate moisture and host emergence was also evident within treatments, as host abundance declined with increasing moisture even among logs with the same position (Fig. 4B). Ant occurrence did not differ significantly between ant-accessible standing and lying deadwood (Fig. 2B) and was itself not associated with host abundance, providing no evidence for a corresponding mediating pathway (Fig. 3).

**Figure 2.**
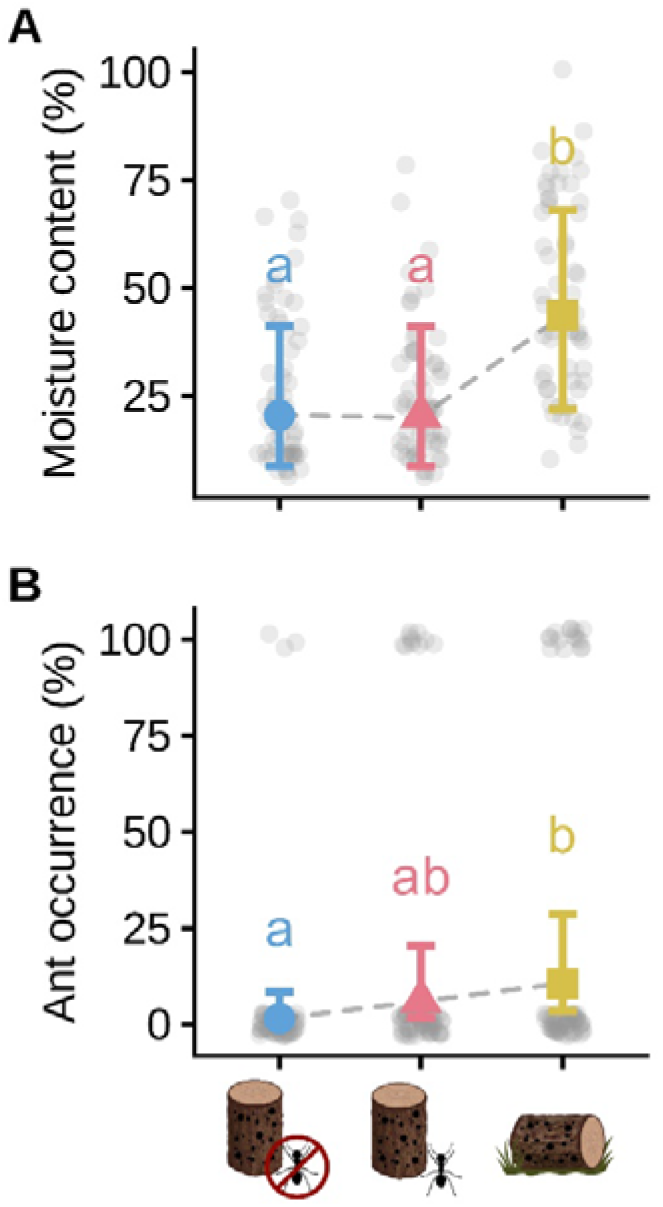
Relationships between standing deadwood with ant exclusion, standing deadwood accessible to ants, and lying deadwood on (A) gravimetric moisture content, and (B) probability of ant occurrence. Measures were taken after deadwood exposure in the field from April 2024 to March 2025 in BEF-China. Symbols represent marginal effects (±95% CI) from GLMMs, points represent individual log traps. Ants breached the exclusion protocol in three (observed) instances. Letters indicate significant differences based on Tukey-adjusted pairwise comparisons.

**Figure 3.**
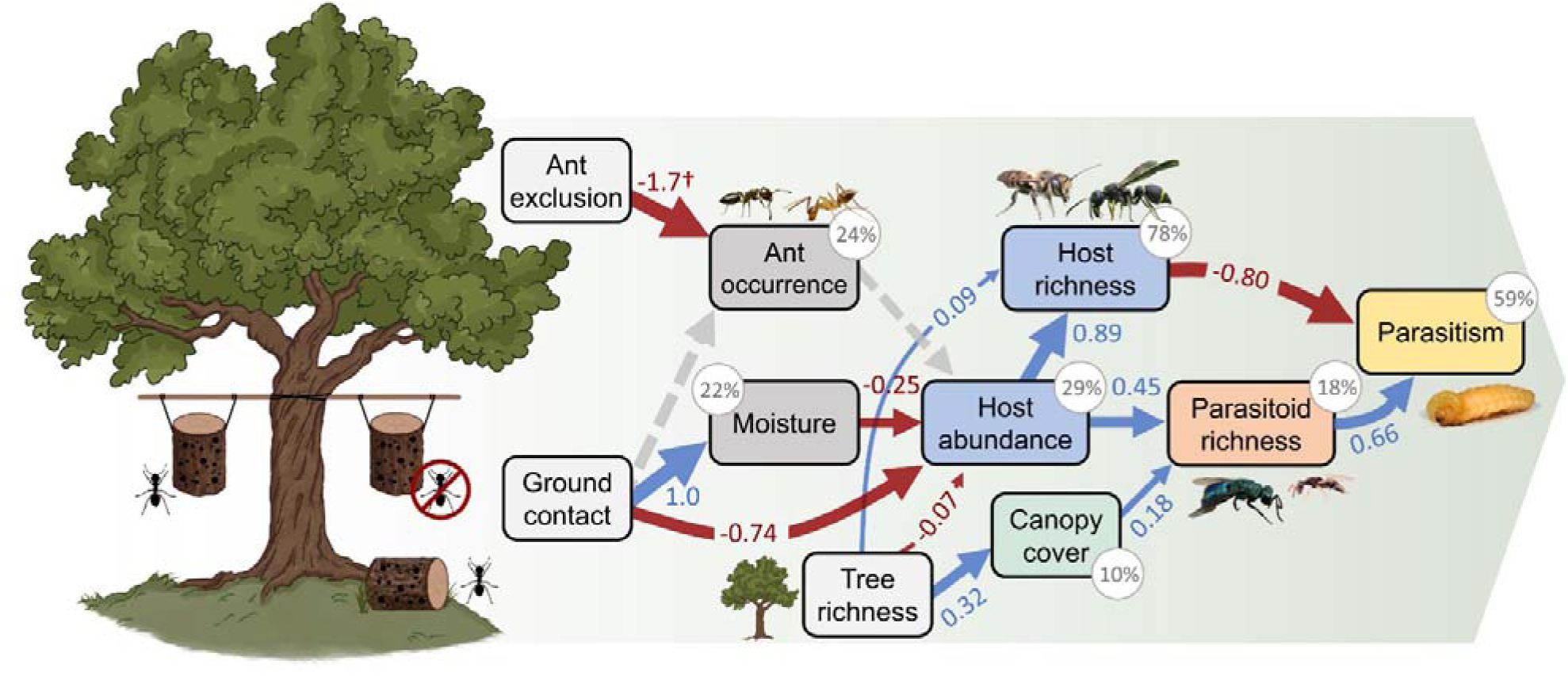
Parsimonious path model of direct and indirect effects of experimentally-manipulated deadwood treatment and tree richness on hosts, parasitoids, and apparent parasitism rate. Component models were fit to response-specific datasets (moisture, 173 traps; ants and hosts, 192 traps; parasitoids and parasitism, 127 traps; canopy cover, 64 plots; see Methods B2). Blue arrows show significant positive effects, red arrows indicate significant negative effects. Path width is scaled according to its standardized coefficient. Grey dashed arrows indicate no effect, showing how observed ant occurrence was not associated with host-parasitoid diversity and parasitism. Other unsupported paths retained in the model are omitted for visual clarity. † indicating log-odds ratio for binomial data (not standardized). Percentage values show the marginal explained variance of endogenous variables. Treatment nodes represent planned contrasts (Methods B2). Standardized coefficients for continuous pathways and *R*² values were derived from a parsimonious standardization model. The path analysis fit the data well (Fisher’s *C* = 15.9, *df* = 20, *p* = 0.72).

**Figure 4.**
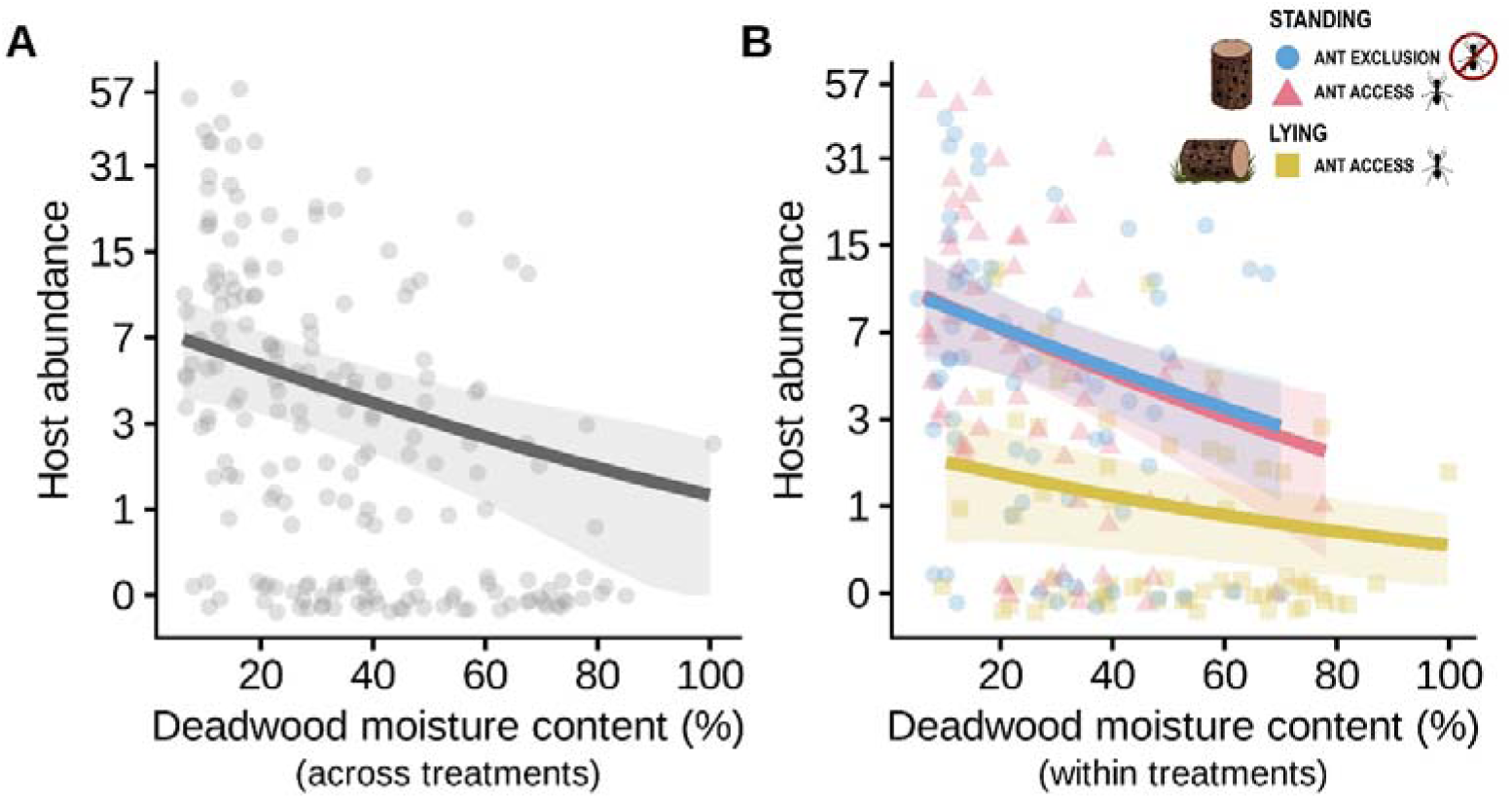
(A) across-treatments and (B) within-treatments marginal associations between deadwood moisture and abundance of cavity-nesting bees and wasps (after accounting for deadwood treatment). Species richness patterns were largely accounted for by variation in abundance. Lines represent marginal effects (±95% CI) from GLMMs. Y-axis is displayed on a log(y + 1) scale for improved visualization, with tick labels shown in the original response units.

Macrohabitat had weaker effects on host communities than microhabitat conditions. Tree species richness showed a weak negative relationship with host abundance (Table C1), but this pattern was strongly influenced by aggregations of the bee *Megachile spissula* Cockerell in several monoculture plots rather than a consistent decline across the richness gradient (Fig. A7). Across mixed-species stands only, host abundance did not vary systematically with tree richness. Host richness increased marginally with tree species richness after accounting for host abundance (Fig. 3, Table C1). However, this association was not supported when richness was standardized to 90% sample coverage or when we used Shannon diversity as response (Table D1). Parasitoid abundance and richness were not directly associated with tree species richness, instead tracking host abundance (Table C1). However, both increased with canopy cover independently of hosts (Fig. 3, Fig. A8, Table C1). Coarse woody debris was uniformly low in the young experimental forest (3.8 ± 0.6 m^3^/ha per plot), and had no detectable effect on hosts, although a weak negative association was observed with parasitoid abundance (Fig. 8, *p* = 0.026). Forest structure did not influence microhabitat conditions: deadwood moisture and ant occurrence were unrelated to tree species richness, canopy cover, or (the latter) to leaf-litter ant occurrence. Instead, deadwood located in valleys generally retained more moisture and was more likely to host ants (Table C1).

Parasitism showed no net response to deadwood position or tree species richness (Fig. 1E), despite strong associations with host and parasitoid diversity. Parasitism increased with parasitoid richness but declined with host richness (Fig. 3). As a result, opposing effects of host and parasitoid diversity largely canceled each other out across treatments. Parasitism only showed an indirect association with canopy cover via direct effects on parasitoid richness (Fig. 3).

Deadwood position sorted host communities into distinct assemblages (r^2^ = 0.07, p < 0.001; Fig. 5A), whereas ant exclusion had no effect. In contrast to abundance patterns, within-treatment moisture variation did not explain additional differences in host community composition. Indicator species analysis identified six species significantly associated with standing deadwood, including the bees *M. spissula* and *Hylaeus sp.*, and the wasps *Anterhynchium flavomarginatum* (Smith), *Pareumenes quadrispinosus* (de Saussure), as well as *Trypoxylon bicolor* (Smith) and *pacificum* Gussakovskij. Only two morphospecies (∼7% of sampled species), *Deuteragenia* sp.2 and *Auplopus* sp.2, were highly associated with lying deadwood. Overall, 18 host species (70%) only occurred in standing deadwood (Fig. 5B). Community dissimilarity between standing and lying deadwood (β_sor_ = 0.58) was largely driven by nestedness (78%), with little species turnover. Tree species richness structured host communities weakly. Plot-level PERMANOVA based on abundance showed no significant effect of tree species richness (r^2^ = 0.029, p = 0.07), whereas the binary analysis indicated a marginal shift in composition (r^2^ = 0.036, p = 0.05), suggesting that tree richness influenced species occurrence rather than their relative abundances.

**Figure 5.**
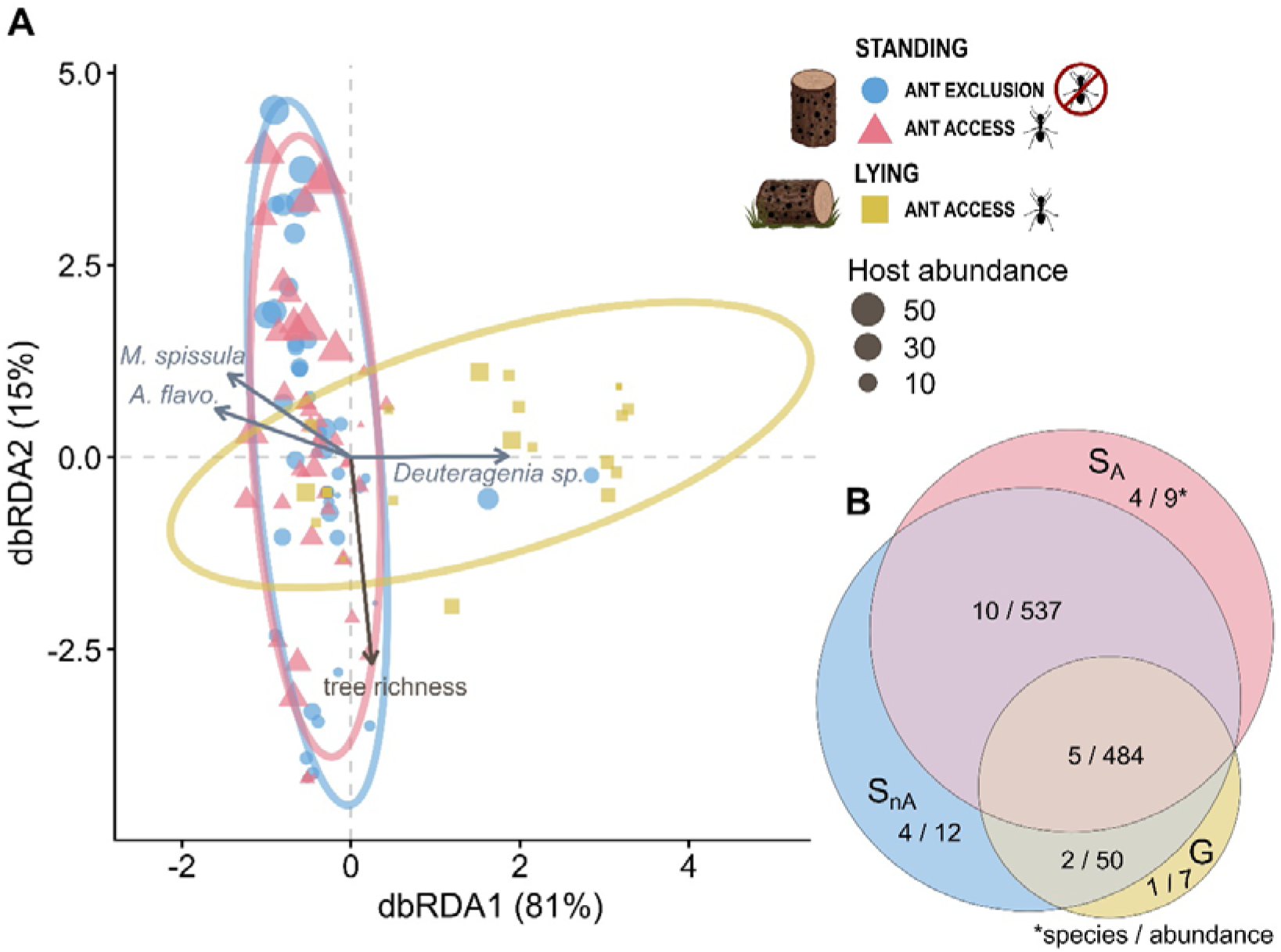
(A) Ordination plot from distance-based redundancy analysis (dbRDA) of deadwood cavity-nesting bee and wasp communities (n = 125, 61 plots). The ordination is constrained by deadwood treatment (standing, standing with ant exclusion, lying) and plot-level tree species richness, based on Bray-Curtis dissimilarities. Axes show the percent constrained variation explained. The black arrow indicates the effect of tree species richness on community composition. Silver arrows indicate host species most strongly associated with the canonical axes, including *Deuteragenia sp.*2 (Pompilidae), *Megachile spissula* (Megachilidae), and *Anterhynchium flavomarginatum* (Vespidae). (B) Euler diagram showing the shared proportion of host species (and summed abundance) between the three deadwood treatments.

## Discussion

While several studies have examined drivers of community assembly in deadwood, far fewer have manipulated habitat characteristics at multiple scales, and large taxonomic and geographic gaps remained (Seibold et al., 2015). Our results provide empirical evidence that deadwood position structures cavity-nesting Hymenoptera communities and consistently identify differences in substrate moisture as an important contributing factor.

Deadwood lying on the forest floor represented a lower-quality nesting substrate for most of the cavity-nesting community, rather than a distinct alternative habitat to standing deadwood. Following our hypotheses, lying deadwood supported taxonomically and numerically reduced communities that were largely nested within those occurring in standing deadwood, suggesting that ground contact acts as a strong environmental constraint. Community patterns across deadwood substrates can be highly taxon-specific (Müller et al., 2020). As non-obligate saproxylics, cavity-nesting Hymenoptera may depend more strongly on the availability and suitability of pre-existing cavities and on favorable microclimatic conditions than intrinsic deadwood traits such as identity, density, or size (Jonsell et al., 2023; Ulyshen, 2018). These insects typically favor dry, warm, and sun-exposed nesting substrates, conditions commonly associated with standing deadwood (Harmon et al., 2011). To our knowledge, only one previous study has directly compared cavity-nesting Hymenoptera communities between standing and lying deadwood and similarly reported higher nesting activity in standing substrates, particularly in elevated sections (Westerfelt et al., 2015). We found reduced parasitoid abundance and diversity in lying compared to standing deadwood. Similar patterns have previously been attributed to lying logs being less accessible to parasitoids (Hedgren, 2007; Ulyshen et al., 2011). Host habitat location typically precedes host location in the sequence of host-searching behavior, indicating that deadwood substrate type may be of particular importance for specialized parasitoid species (Vinson, 1976; Ulyshen, 2018). However, reduced parasitoid diversity in lying deadwood was fully mediated by a reduced availability of hosts. This indicates that the detectability and accessibility of hosts mainly decreased with host abundance, rather than through diminished exposure of deadwood due to contact with the ground (Dynesius et al., 2010).

As expected, our results identify increased substrate moisture as an important variable associated with reduced host emergence. Lying deadwood was wetter and supported fewer hosts, while host abundance also declined with increasing moisture among deadwood objects in the same position. Elevated moisture can impair Hymenoptera larval development and survival, for example through the promotion of fungal growth within brood cells (Stephen, 1965; Flores et al., 1996). Such constraints likely reduce the suitability of wetter deadwood as nesting substrate. Although substrate humidity is an expected limiting factor for cavity-nesting Hymenoptera, direct experimental evidence remained scarce. Our results suggest that moisture gradients may constrain these communities at two complementary scales: broad differences between deadwood types define general suitability thresholds, while finer variation within substrates further modulates host abundance. Future studies could build on these findings by tracking substrate moisture throughout nesting and brood development while quantifying fungal activity and wood deterioration, thereby distinguishing correlated effects of ground contact from processes downstream of moisture.

Moisture mediated part of the effect of deadwood position on community size, but not community composition. A few species (∼7% of total) were associated with lying deadwood, suggesting some potential variation in moisture tolerance between taxa. However, most were absent or occurred less frequently in wetter substrates. Moisture was thus associated primarily with lower abundance and diversity of overwintering hosts, which could reflect reduced substrate receptivity, brood cell construction, brood survival, or a combination of these processes. Compositional differences between standing and lying deadwood were therefore consistent with a nested species loss rather than a turnover towards moisture-tolerant taxa.

Contrary to expectations, we detected no difference in host or parasitoid communities between ant-exclusion and ant-accessible standing deadwood. Ants are considered dominant antagonists in deadwood and can limit arthropod communities and associated ecosystem functions (Law et al., 2019; Wu et al., 2021), although neutral or positive interactions are also possible (Ulyshen et al., 2020). In our system, neither ant exclusion from standing deadwood nor observed ant occurrence in standing and lying deadwood was associated with host abundance or diversity. This differs from results from another subtropical region, where ant exclusion from reed-filled trap nests increased nesting frequency and revealed antagonistic relationships that were strongest in autumn (Miyano & Yamaguchi, 2001). Under the conditions captured by our experiment, our results provide little evidence that ants are a major driver of cavity-nesting Hymenoptera community assembly or trophic interactions, particularly in standing deadwood. However, most ant-accessible deadwood traps (∼80%) had no ants at final collection, resulting in a weak realized treatment contrast (Fig.2B) and limiting power to detect ecological effects. Remaining observations generally involved few individuals rather than colonial aggregations, suggesting limited use of the experimental substrate. Ant nesting in deadwood generally increases with forest age and wood decay stage (Boucher et al., 2015; Tanaka et al., 2023), suggesting that the young experimental forest or relatively fresh logs may have provided suboptimal colonizing conditions. Critically, as ant occurrence was recorded only once logs were removed in spring, we could not capture seasonal foraging, predation, or deterrence throughout the annual cycle (Kass et al., 2023). The absence of ants at final collection therefore does not rule out earlier transient activity or, although the sticky resin glue appeared largely effective, additional breaching of barriers earlier in the experiment. Finally, since ants were not excluded from lying deadwood, any unmeasured ant activity associated with ground contact could not be disentangled from the position effect, and may thus have accounted for part of the substantial residual difference not explained by the moisture pathway (Fig. 3). Future studies may need to apply broad (plot-scale) ant suppression in order to study ant exclusion effects in lying deadwood without also disrupting the study organisms, and monitor ant activity at regular intervals throughout the field experiment.

Beyond the strong effects of deadwood position, macrohabitat gradients acted as comparatively weak constraints for bee and wasp communities, following our original hypothesis. Host richness and community composition responded weakly to tree species richness, whereas host abundance did not vary consistently across mixed-species stands and neither canopy cover nor coarse woody debris had a detectable effect. Moreover, the positive tree-host richness association was absent for coverage-standardized richness and Shannon diversity, suggesting that it primarily reflected the occurrence of rare species rather than changes in the abundance or evenness of dominant taxa. This was also consistent with plot-level community analyses showing a slightly higher association between tree richness and incidence-based compositional variation compared to abundance-based variation. Our results contrast with studies using conventional reed trap nests, which have reported relationships with canopy cover, stand volume, structural complexity, tree composition, and the availability of standing deadwood (Fornoff et al., 2021; Rappa et al., 2023, 2024; Wang et al., 2025; Martini et al., 2026). This may suggest that standardized nesting aids may make variations in surrounding forest structure more apparent. Instead, strong variations in substrate conditions created by deadwood position may have constrained nest establishment or brood survival before macrohabitat resources became limiting. Our results therefore do not imply that forest context is unimportant, but suggest that its influence may be secondary when local nesting conditions impose a strong environmental filter. Moreover, although mixed-effect models accommodate unbalanced designs, declining replication along the tree-richness gradient may have limited our ability to detect modest or nonlinear relationships of the kind reported in larger, more balanced studies (Martini et al., 2026).

Parasitoids showed a somewhat stronger response than hosts to forest structure. In particular, canopy cover was positively associated with parasitoid abundance and diversity and, consequently, with parasitism (although the latter indirect relationship was not significant; *p* = 0.06). Because canopy cover increased with tree species richness in the experimental forest, this pattern suggests that tree diversity indirectly supported parasitoids through structural habitat changes. Such responses are consistent with expectations that natural enemies benefit directly from increased habitat complexity and structural heterogeneity (Staab & Schuldt, 2020). Our results indicate that parasitoids responded more strongly than their hosts to forest structure, highlighting how macrohabitat conditions can differentially constrain trophic levels within saproxylic communities. Coarse woody debris volume also showed little influence, with no association to hosts and a weak negative correlation with parasitoids. While positive relationships are expected between deadwood volume and cavity-nesting Hymenoptera and their parasitoids (Rappa et al., 2023, 2024), the uniformly low CWD values in the young experimental forest were well below thresholds typically associated with strong responses of saproxylic communities (Müller & Bütler, 2010). Increased structural complexity from CWD could theoretically reduce parasitoid search efficiency (Gingras et al., 2002), however our results likely reflect unmeasured heterogeneity of parasitoid abundance across the restricted CWD gradient, rather than a true ecological effect. As the forest matures and deadwood increases, CWD may become a more robust driver of host and parasitoid communities.

Although deadwood position structured host and parasitoid communities, apparent parasitism did not differ significantly among treatments. Instead, it increased with parasitoid richness and declined with host richness, consistent with complementarity or selection among enemies and dilution through reduced encounters with suitable hosts (Tylianakis et al., 2006; Civitello et al., 2015). These opposing relationships may partly buffer parasitism against environmentally driven changes in community structure, illustrating the value of considering multiple trophic levels when linking biodiversity to ecosystem functioning.

## Conclusions

Our study identifies substrate moisture as an important factor limiting bee and wasp communities nesting in deadwood cavities. In contrast, forest characteristics such as tree diversity, canopy cover, and coarse woody debris played minor roles – also in comparison to conventional trap nest literature – while our exclusion treatment and observations provided little evidence of negative ant interactions. Our results provide a candidate explanation for previously-reported positive associations between cavity-nesting bees, wasps, and their parasitoids and standing deadwood in forests. Because standing deadwood is typically scarce in managed forests (Wisdom & Bate, 2007), our findings support a rationale for retaining or creating standing deadwood structures and, more broadly, promoting dry and elevated nesting substrates for cavity-nesting bees and wasps in conservation and restoration measures. Lying deadwood remains an essential component nonetheless, as some species show clear associations with substrates on the ground and many other saproxylic organisms depend on this microhabitat.

## Acknowledgements

We gratefully acknowledge the BEF-China platform and the Zhejiang Qianjiangyuan Forest Biodiversity National Observation and Research Station (QForDiv). Special thanks to Keping Ma for initiating the BEF-China platform, and Bo Yang for his efforts in maintaining the research station. We thank the local and student helpers for assistance in the field and laboratory. We are grateful to Olivia Scheel for her original illustrations.

## Data availability statement

Data and code for statistical analyses is archived on Zenodo and maintained in a public GitHub repository, available at: https://doi.org/10.5281/zenodo.22177977. Data is also available on Fighsare at: https://doi.org/10.6084/m9.figshare.31979481.v2

## Declaration of generative AI in scientific writing

During the preparation of this work the authors used Open AI’s ChatGPT in order to improve code for data analysis and readability of manuscript text. During and after using this tool, the authors reviewed and edited the content as needed and take full responsibility for the contents of the publication.

## Author contributions

Conceptualization by Massimo Martini, Matteo Dadda, Felix Fornoff, Heike Feldhaar, Chao-Dong Zhu, Alexandra-Maria Klein; Investigation by Massimo Martini, Matteo Dadda, Joshua Spitz; Funding acquisition by Alexandra-Maria Klein; Data curation by Massimo Martini, Matteo Dadda, Finn Rehling, Joshua Spitz; Formal analysis by Massimo Martini, Matteo Dadda, Felix Fornoff, Finn Rehling; Validation by Massimo Martini, Finn Rehling; Visualization by Massimo Martini, Felix Fornoff, Finn Rehling. All authors contributed critically to the drafts and gave final approval for publication.

## Appendix A. Supplementary Figures and Tables

**Figure A1.**
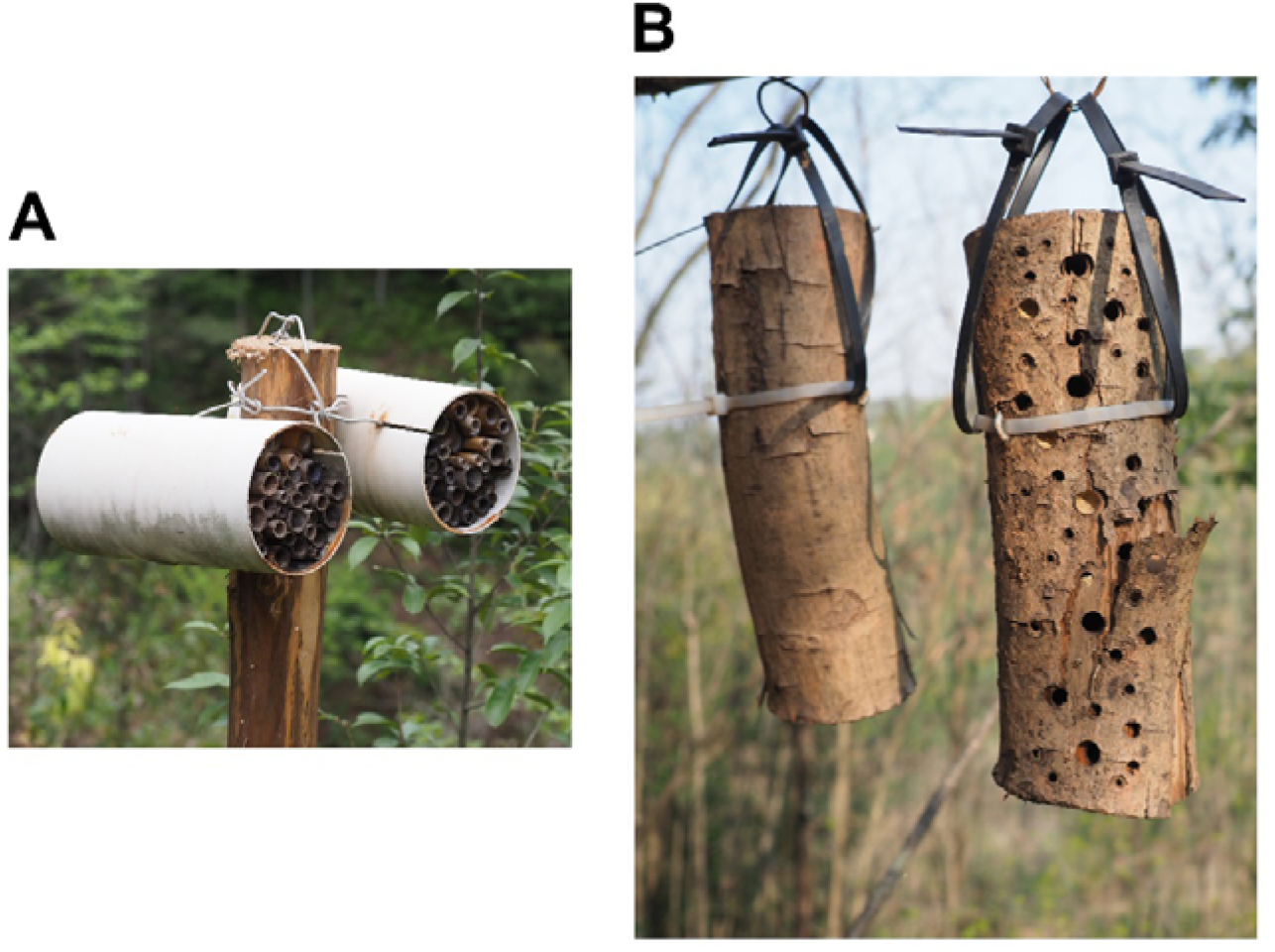
(A) Standard trap nest filled with hollow reeds. (B) Our new deadwood trap for sampling deadwood-cavity-nesting bees and wasps and their natural enemies. Here visualized is a trap mimicking standing (or suspended) deadwood. Traps consist of ∼10 x ∼30 cm logs (diameter & length) of *Schima superba,* felled in October 2023 and drilled with cavities of 3 – 12 mm diameter (see table S1). They were deployed and left untouched in the field from April 2024 to March 2025. After removal, the deadwood traps were kept in fine-mesh bags until late August to collect all emerging insects. Adjacent to all deadwood traps were logs without drilled cavities, used to measure the gravimetric water content (GMC) after removal.

**Figure A2.**
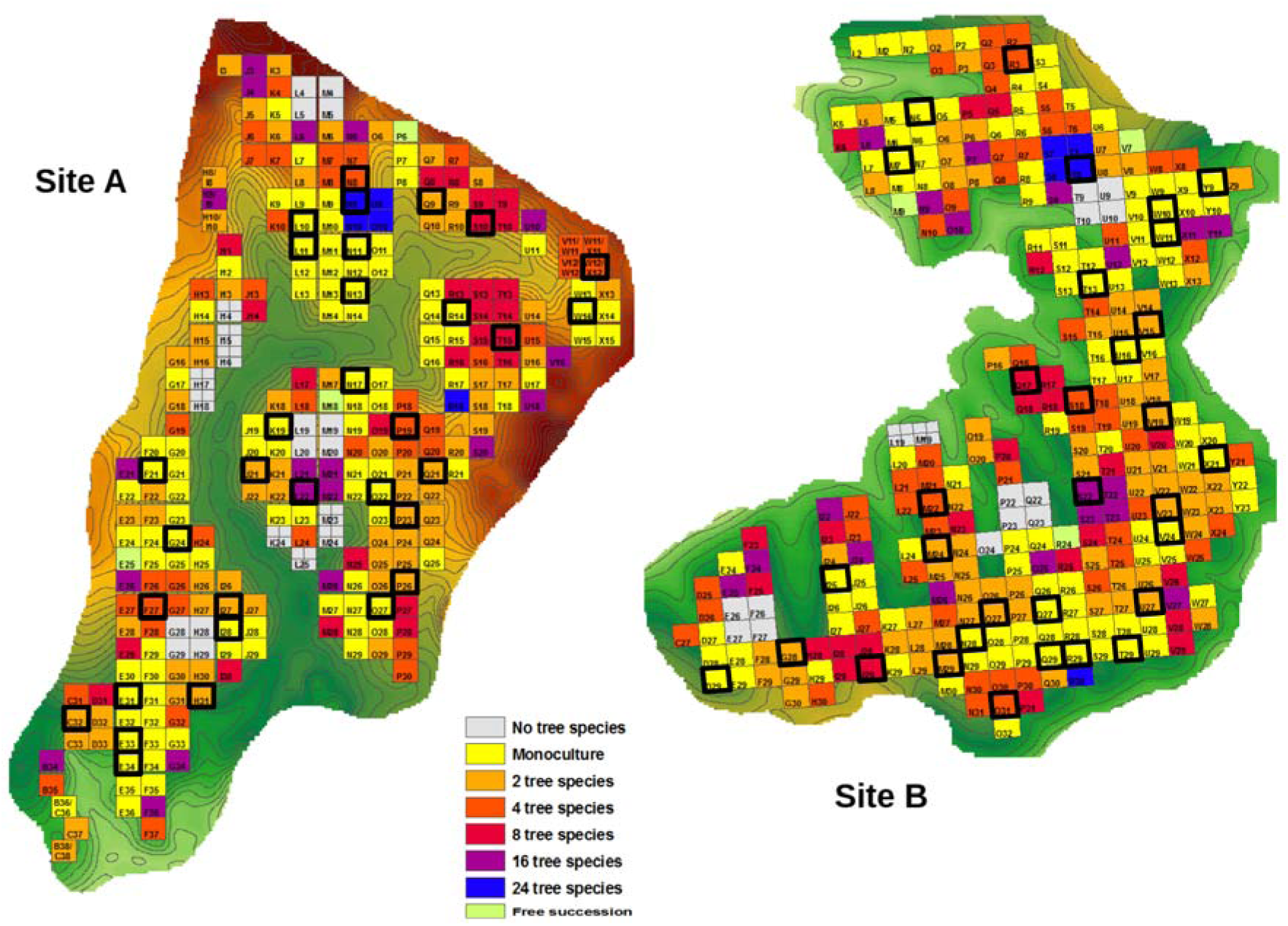
Topographic maps of the BEF-China experimental forest sites. Site A (left) and B (right) occupy approximately 20 ha each and were planted in 2009 and 2010, respectively. Overall, 566 study plots of 1/15 ha were established by planting seedlings from a pool of 40 native tree species. Highlighted in black are the 64 study plots where deadwood traps were placed.

**Figure A3.**
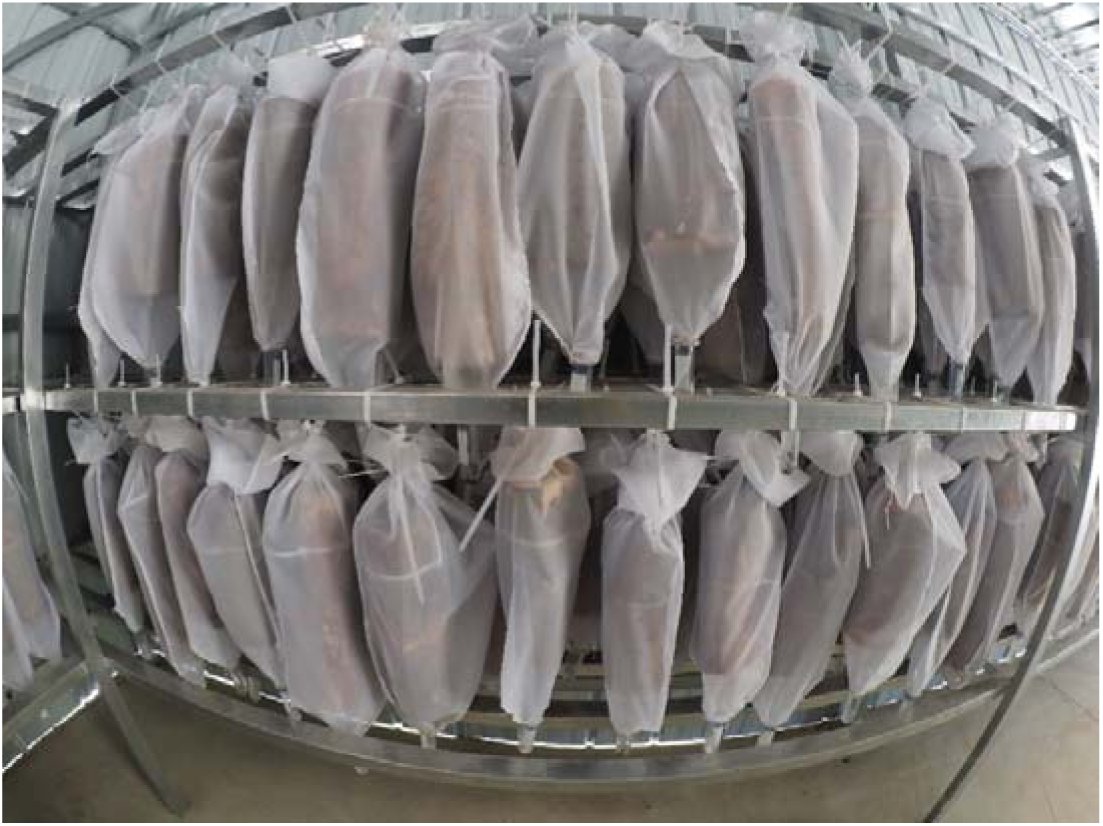
Suspended deadwood traps wrapped in fine-mesh nylon bags with collection vials at the bottom for emerging insects. The logs were placed in an outdoor shed immediately after removal from the field, protected from rain but exposed to the natural temperature fluctuations. Emergence of overwintering insects lasted for five months, from March to August 2025.

**Figure A4.**
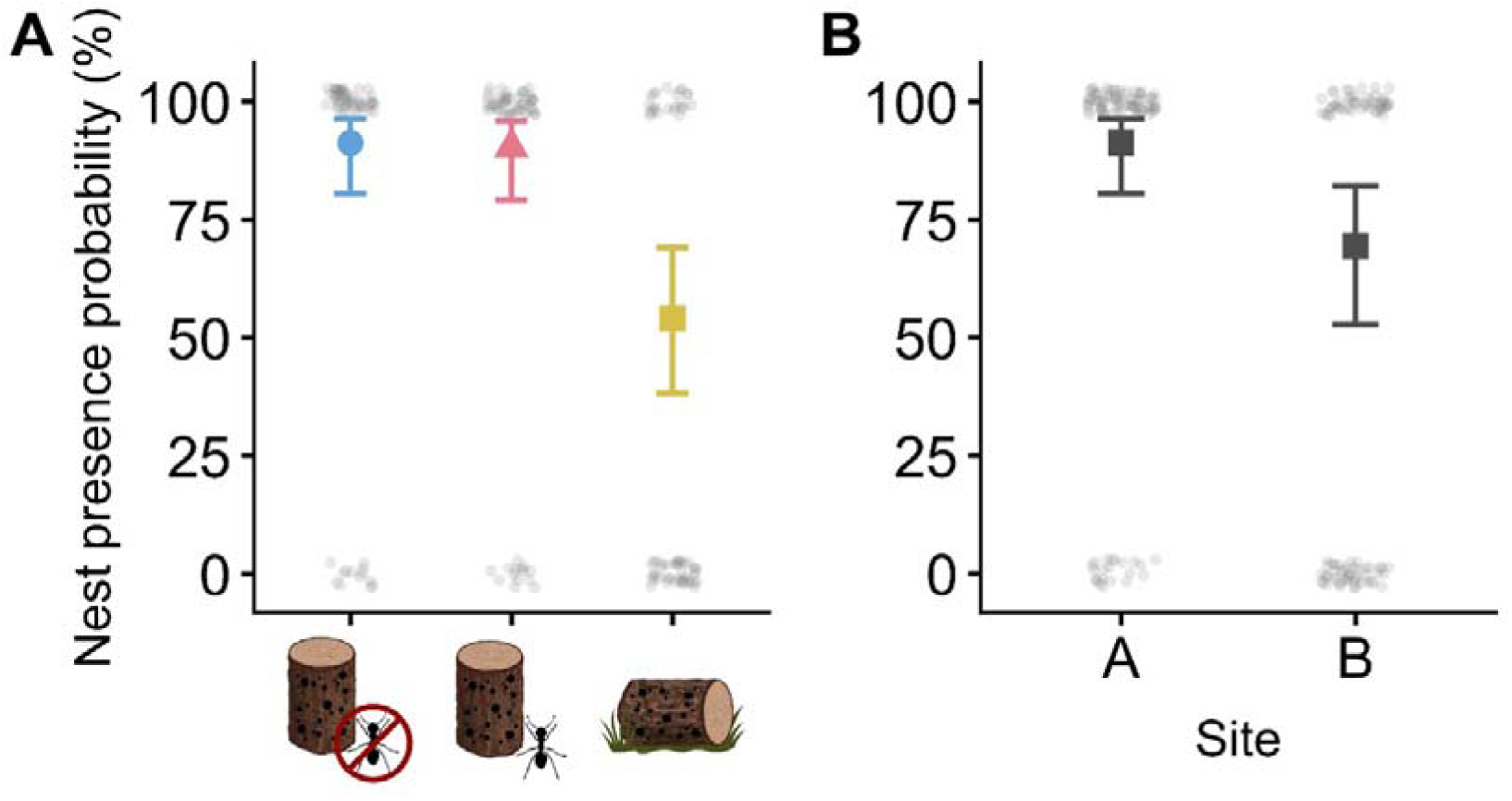
Probability of nest occurrence (±95% CI) in deadwood traps across (A) deadwood treatments, and (B) forest sites. Lying deadwood, and any deadwood in site B on average were less likely of housing overwintering host nests. Results are from GLMM model on binary host presence-absence data.

**Figure A5.**
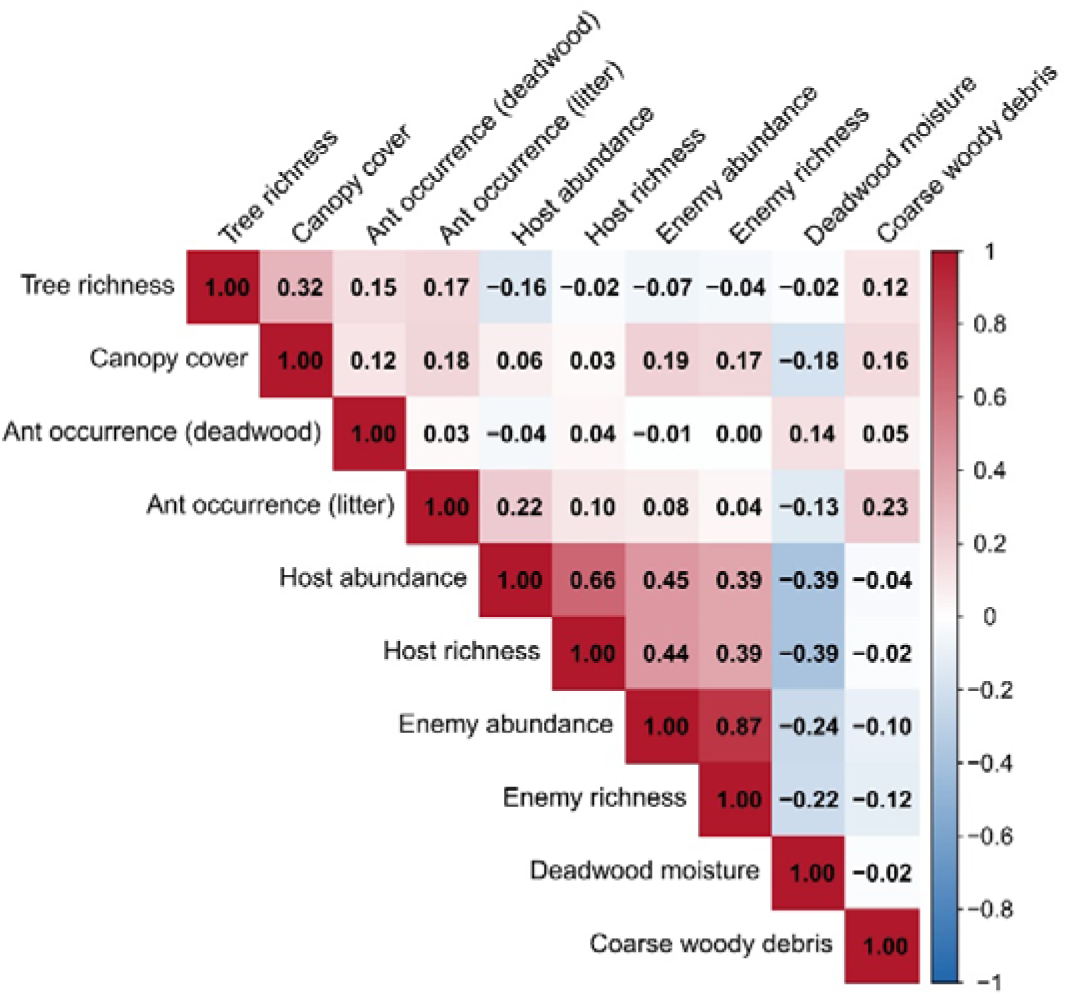
Pairwise rank correlations among log, plot, forest, host, and enemy variables. The pairwise correlation matrix gives coefficients (Spearman’s ρ) between variables used in path analyses and mixed-effects models. Red indicates positive and blue negative associations; color saturation reflects correlation strength.

**Figure A6.**
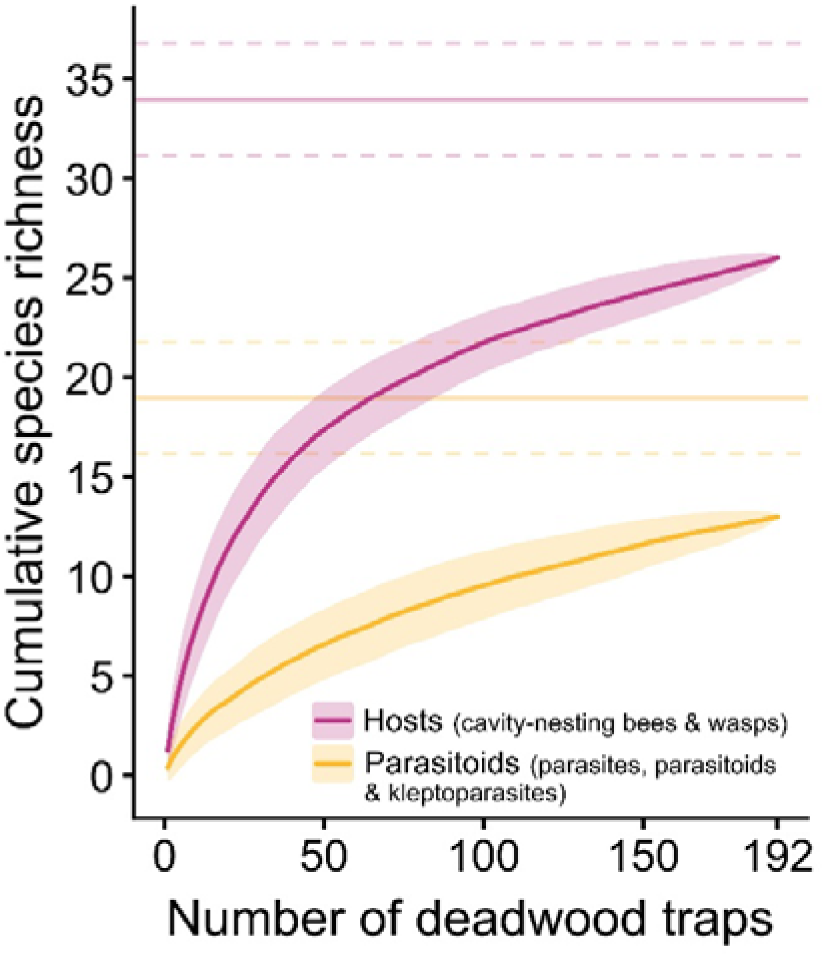
Species accumulation curves for solitary cavity-nesting hosts and their parasitoids. Calculated based on total number of deadwood log traps exposed in the field. Shown are the observed number of species (solid lines), the ±1 SD of the accumulation curves from 1,000 random permutations (shading), and the expected number of species ± SE based on jack1 estimators (solid and dashed horizontal lines, respectively). Approximately 76% (26 species) of the total expected host species, and 68% (13 species) of the total expected natural enemy species were sampled.

**Figure A7.**
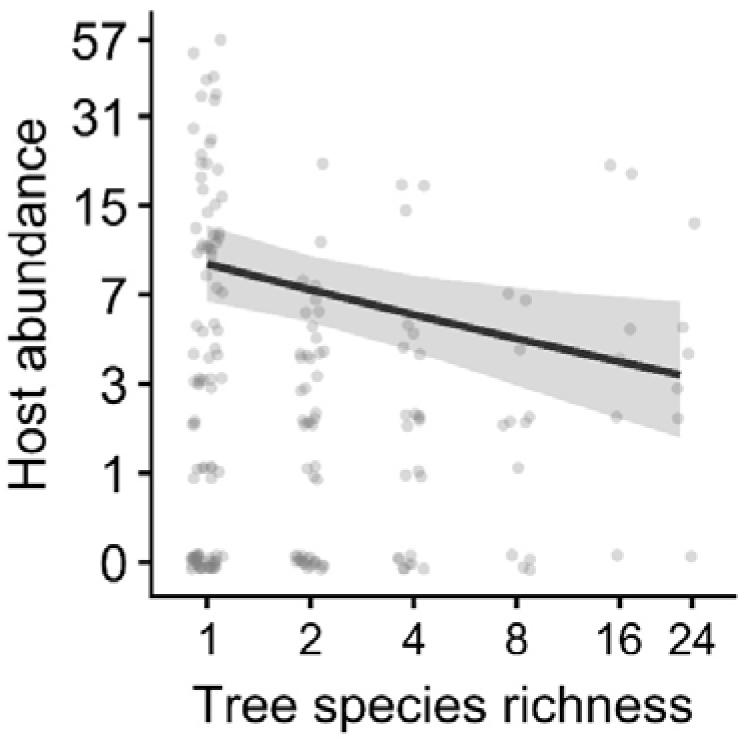
Effect of tree species richness on deadwood cavity-nesting Hymenoptera abundance. Line shows predicted marginal effect from GLMM model (± 95% CI). The negative correlation was no longer significant when monocultures were excluded from the model.

**Figure A8.**
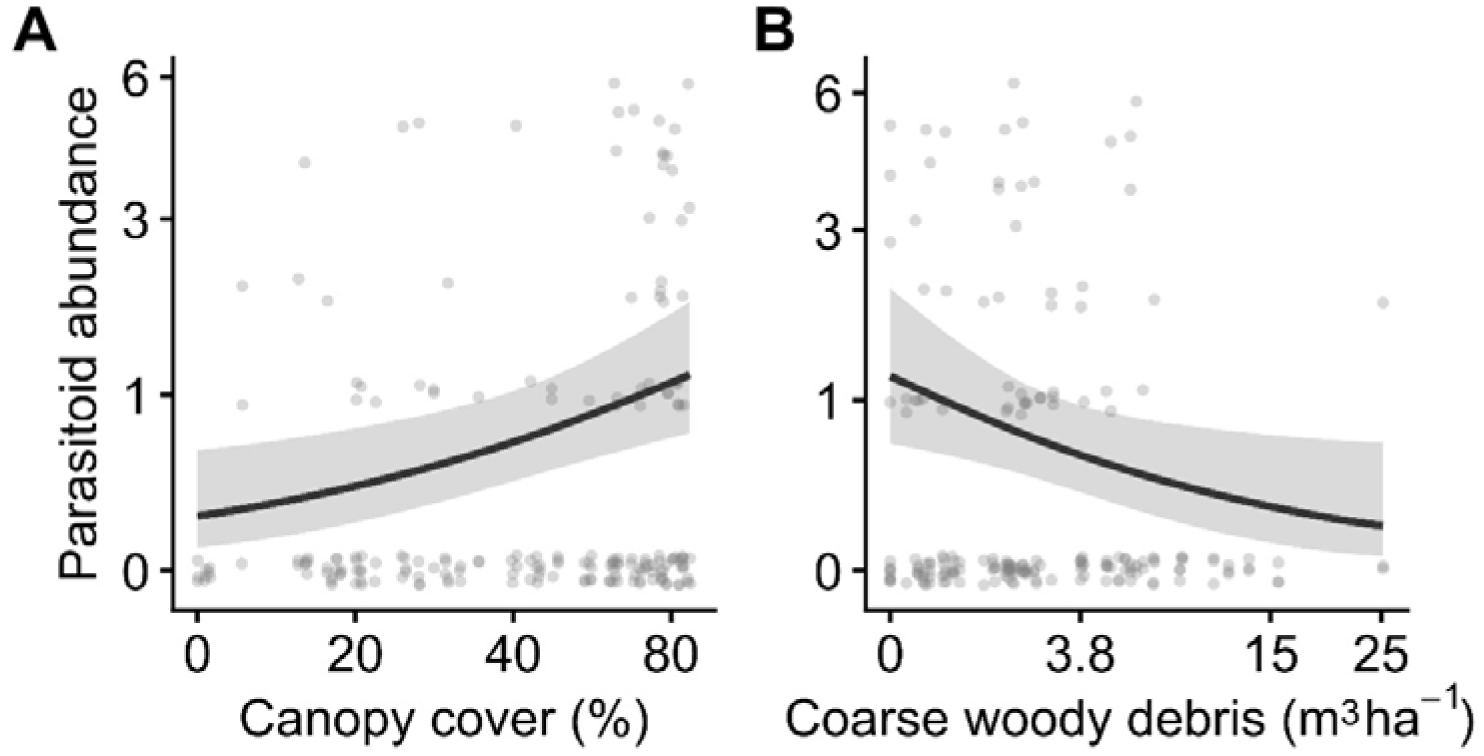
Effect of canopy cover (A) and coarse woody debris (CWD, panel B) on parasitoid abundance in deadwood traps. Lines represent marginal effects from GLMM (± 95% CI).

**Table A1.** Distribution of pre-drilled cavity diameters and depths per deadwood trap. This follows the distribution of reed diameters selected by the same cavity-nesting Hymenoptera community reported from previous studies.

| Diameter (mm) | Depth (mm) | Number (per trap) |
| --- | --- | --- |
| 3 | 33 | 5 |
| 4 | 43 | 7 |
| 5 | 52 | 10 |
| 6 | 57 | 6 |
| 8 | 75 | 6 |
| 10 | 87 | 5 |
| 12 | 95 | 5 |

**Table A2.**
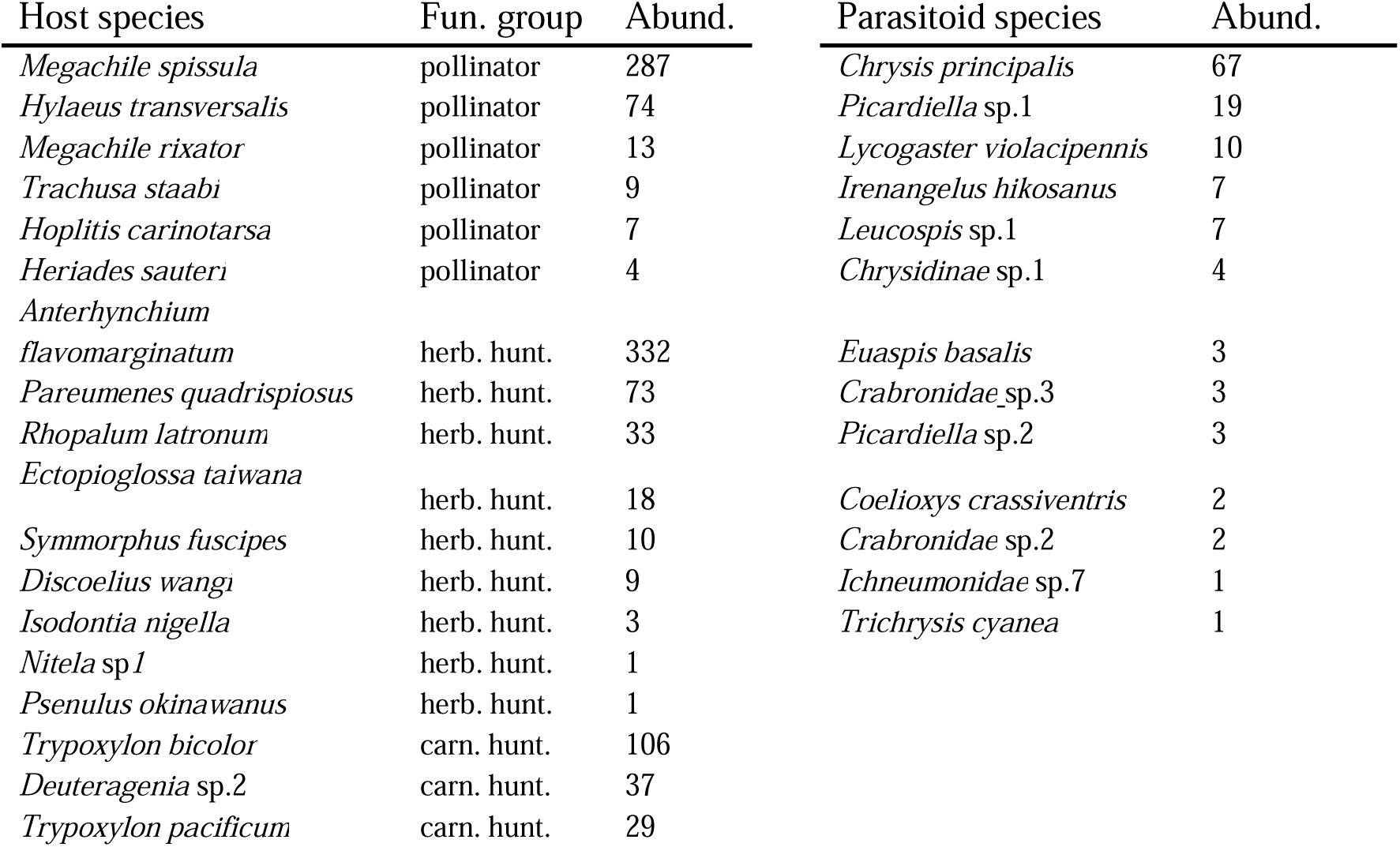
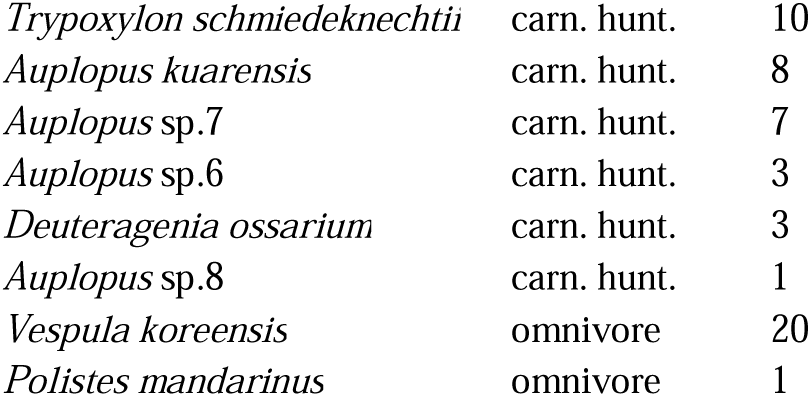
Species list and abundance (number of emerged adults) of deadwood cavity-nesting host and parasitoid species (parasites, parasitoids and kleptoparasites) observed in deadwood traps from BEF-China (site A and B). Species are ordered by decreasing abundance, hosts additionally by functional group (pollen-collecting bees, herbivore-hunting or carnivore-hunting wasp).

## Appendix B. Supplementary Methods

### Methods B1. Data imputation

A small proportion of predictor variables contained missing values: deadwood moisture content was absent for 19 logs (9.9% of log-level observations), coarse woody debris volume for 4 plots (6.25%), and litter ant occurrence for 3 plots (4.7%). Missing values were distributed across different plots. Listwise deletion would have removed 25 logs (13% of the total). We therefore imputed missing values using GLMM model-based predictions fitted to complete cases, a principled approach that leverages the correlation structure among predictors while preserving sample size (Nakagawa and Freckleton, 2008; Van Buuren, 2018). Deadwood moisture was imputed using a beta regression mixed model including treatment and plot-level structural variables, topographic covariates, and random intercepts for plot nested within site; missing values were replaced with fitted values on the response scale. CWD volume and leaf litter ant occurrence were each imputed at the plot level using Gaussian mixed models with forest and topographic predictors and a random intercept for site; missing values were drawn stochastically from a normal distribution centred on model predictions with standard deviation equal to the model residual standard deviation, preserving uncertainty. Imputed values were truncated at zero and ant occurrence values were rounded to the nearest integer. All imputed variables were used exclusively as predictors in downstream models and never as response variables to avoid circular inference.

### Methods B2. Path analyses

#### Data preparation

Path analyses were conducted at the deadwood-trap level using 192 traps from 64 plots. Tree richness was log2-transformed and host abundance was log(y + 1)-transformed. Apparent parasitism was calculated as *P*/(*H* + *P*), where *P* was parasitoid abundance and *H* successfully emerged host abundance, and subsequently bounded between 0 and 1 for beta regression. Host and ant components used all 192 traps. Parasitoid richness and apparent parasitism used the the 127 traps with evidence of host brood, defined as successful host emergence or parasitoid emergence. The moisture component model used the 173 traps with measured moisture. Canopy cover was represented once per plot (*n* = 64) to avoid pseudoreplication. The pSEM was supplied the full 192-row dataset for tests of directed separation. The full and parsimonious response-specific component models were specified as shown in the following table:

| Response | Sample | Distribution and additional structure | Random effect | Full-model predictors | Parsimonious-model predictors |
| --- | --- | --- | --- | --- | --- |
| Apparent parasitism | 127 | Beta, logit link | site/plot | treatment, moisture, ant occurrence, parasitoid richness, host richness, tree richness, canopy cover | treatment, moisture, parasitoid richness, host richness, canopy cover |
| Parasitoid richness | 127 | Poisson, log link | none | treatment, host richness, host abundance, ant occurrence, moisture, tree richness, canopy cover | treatment, host abundance, moisture, canopy cover |
| Host richness | 192 | Generalized Poisson, log link | plot | treatment, moisture, tree richness, canopy cover, host abundance | tree richness, canopy cover, host abundance |
| Host abundance | 192 | Zero-inflated Gamma, log link; zero-inflation: site + treatment | site/plot | treatment, tree richness, moisture, ant occurrence, canopy cover | treatment, tree richness, moisture, canopy cover |
| Moisture | 173 | Beta, logit link; dispersion: site + treatment | site/plot | treatment, tree richness, canopy cover | unchanged |
| Ant occurrence | 192 | Binomial, logit link | site/plot | treatment, tree richness, canopy cover | unchanged |
| Canopy cover | 64 | Gaussian, identity link | site | tree richness | unchanged |

*Notes. Host abundance was log(y + 1)-transformed prior to all path analyses. Random effects were removed if near-zero variance caused model convergence failure.*

Parasitoid abundance was excluded from path analyses due to its strong redundancy with parasitoid richness (*r* = 0.87). The maximal model retained every hypothesized pathway. The parsimonious model was derived by sequentially removing selected unsupported pathways while monitoring log-likelihood-based piecewise AIC and Fisher’s C; it was retained as a simplified representation rather than as a replacement for the full model. Both structures and their complete coefficient tables are reported in Appendix B.

#### Matched Gaussian-standardization models

While piecewise SEM can automatically calculate standardized coefficients for some generalized models, this is not supported for all families used in our response-specific components, including beta, generalized-Poisson, and zero-inflated Gamma models. We therefore fitted a matched standardization model for each full and parsimonious response-specific pSEM. Each standardization model matched the directed pathways and correlated errors of its original counterpart. Continuous nodes were transformed where necessary and standardized once using their designated reference populations before fitting Gaussian identity-link components; binary ant occurrence remained binomial. Apparent parasitism was transformed using the empirical logit qlogis (*P* + 0.5)/(*H* + *P* + 1), parasitoid and host richness were square-root transformed, host abundance was log1p-transformed, moisture was log-transformed, tree richness retained its log2 transformation, and canopy cover remained on its proportional scale. Estimates from the matched standardization models were interpreted as standardized coefficients only for continuous-to-continuous pathways. Treatment effects were represented by pre-specified contrasts, while paths involving binary ant occurrence were interpreted as binary-predictor effects rather than standardized continuous slopes. Pathway *p* values and other inferential statistics were obtained from the corresponding response-specific pSEMs.

#### Treatment contrasts

For the parsimonious models, we calculated four a-priori treatment contrasts. With treatment levels ordered: a. standing with ant exclusion, b. standing, and c. lying; ground-contact deadwood was compared with the average of both standing treatments for host abundance and moisture using weights (−0.5, −0.5, 1); ant exclusion was compared with the average of both ant-accessible treatments for ant occurrence using (1, −0.5, −0.5); and ground-contact deadwood was compared directly with ant-access standing deadwood for ant occurrence using (0, −1, 1). The same contrasts were fitted to the response-specific and matched Gaussian components. Response-specific contrasts supplied inferential statistics, while Gaussian contrasts supplied estimates for the standardized SEM figure; no multiplicity adjustment was applied to these prespecified comparisons. Supplementary coefficient tables present response-specific and Gaussian estimates side by side, but SEs, test statistics, degrees of freedom, *p-*values, and all statements about pathway support come from the response-specific pSEMs.

