## Appendix C for "Moisture limits the diversity of nesting bees, wasps, and parasitoids in lying and standing deadwood"

### Appendix C. Statistical model results

Main statistical model results for the manuscript: "Moisture limits the diversity of nesting bees, wasps, and parasitoids in lying and standing deadwood"

Massimo Martini, Matteo Dadda, Felix Fornoff, Heike Feldhaar, Arong Luo, Finn Rehling, Joshua E. Spitz, Michael Staab, Simon Thorn, Chao-Dong Zhu, Alexandra-Maria Klein

C1. Main GLMM results (p. 1-4)

C2. PERMANOVA results (p. 5)

C3. Structural equation model results (p. 6-7)

#### C1. Main GLMM results

Main model coefficient table

| Component | Predictor | Estimate | SE | Test statistic | P | 95% CI |
|---|---|---|---|---|---|---|
| **Moisture content — beta (logit link); n = 173; logLik = 159.539; AIC = -293.078** | | | | | | |
| Conditional | Intercept | -1.357 | 0.511 | -2.656 | **0.008** | -2.358, -0.356 |
| Conditional | Treatment: S[A] vs S[nA] | -0.003 | 0.068 | -0.051 | 0.960 | -0.136, 0.129 |
| Conditional | Treatment: G vs S[nA] | 1.099 | 0.102 | 10.730 | **<0.001** | 0.899, 1.300 |
| Conditional | Tree richness | -0.058 | 0.057 | -1.023 | 0.306 | -0.170, 0.054 |
| Conditional | Canopy cover | 0.053 | 0.061 | 0.863 | 0.388 | -0.067, 0.172 |
| Conditional | Geomorphology: category 2 vs 1 | 0.292 | 0.121 | 2.408 | **0.016** | 0.054, 0.530 |
| Conditional | Geomorphology: category 3 vs 1 | 0.649 | 0.149 | 4.353 | **<0.001** | 0.357, 0.941 |
| Random effects | plot_id:site: (Intercept) | 0.308 | | | | |
| Random effects | site: (Intercept) | 0.709 | | | | |
| **Ant occurrence — binomial (logit link); n = 192; logLik = -68.469; AIC = 156.937** | | | | | | |
| Conditional | Intercept | -4.194 | 0.922 | -4.547 | **<0.001** | -6.002, -2.386 |
| Conditional | Treatment: S[A] vs S[nA] | 1.474 | 0.728 | 2.025 | **0.043** | 0.047, 2.900 |
| Conditional | Treatment: G vs S[nA] | 2.063 | 0.722 | 2.857 | **0.004** | 0.648, 3.478 |
| Conditional | Tree richness | 0.444 | 0.255 | 1.743 | 0.081 | -0.055, 0.943 |
| Conditional | Plot-level ant occurrence | -0.093 | 0.267 | -0.348 | 0.728 | -0.617, 0.431 |
| Conditional | Canopy cover | 0.176 | 0.284 | 0.619 | 0.536 | -0.381, 0.732 |
| Conditional | Geomorphology: category 2 vs 1 | 0.858 | 0.644 | 1.333 | 0.183 | -0.404, 2.120 |
| Conditional | Geomorphology: category 3 vs 1 | 1.666 | 0.720 | 2.314 | **0.021** | 0.255, 3.077 |
| Random effects | plot_id:site: (Intercept) | 0.685 | | | | |
| Random effects | site: (Intercept) | 0.000 | | | | |
| **Host occurrence — binomial (logit link); n = 192; logLik = -97.258; AIC = 206.516** | | | | | | |
| Conditional | Intercept | 2.187 | 0.456 | 4.799 | **<0.001** | 1.294, 3.080 |
| Conditional | Treatment: S[A] vs S[nA] | 0.067 | 0.465 | 0.144 | 0.886 | -0.844, 0.978 |
| Conditional | Treatment: G vs S[nA] | -2.200 | 0.488 | -4.511 | **<0.001** | -3.156, -1.244 |
| Conditional | Site: B vs A | -1.379 | 0.395 | -3.492 | **<0.001** | -2.153, -0.605 |
| Conditional | Ant presence | 0.513 | 0.533 | 0.961 | 0.336 | -0.533, 1.558 |
| Random effects | plot_id: (Intercept) | 0.179 | | | | |
| **Host abundance — nbinom2 (log link); n = 192; logLik = -479.762; AIC = 989.523** | | | | | | |
| Conditional | Intercept | 2.000 | 0.148 | 13.494 | **<0.001** | 1.710, 2.291 |
| Conditional | Treatment: S[A] vs S[nA] | -0.015 | 0.173 | -0.087 | 0.930 | -0.355, 0.325 |
| Conditional | Treatment: G vs S[nA] | -1.279 | 0.273 | -4.686 | **<0.001** | -1.814, -0.744 |
| Conditional | Tree richness | -0.284 | 0.108 | -2.631 | **0.009** | -0.496, -0.072 |
| Conditional | Moisture content (within treatments) | -0.331 | 0.109 | -3.043 | **0.002** | -0.544, -0.118 |
| Conditional | Ant presence | 0.189 | 0.267 | 0.708 | 0.479 | -0.334, 0.713 |

| Component | Predictor | Estimate | SE | Test statistic | P | 95% CI |
|---|---|---|---|---|---|---|
| **Host abundance — nbinom2 (log link); n = 192; logLik = -479.762; AIC = 989.523 (continued)** | | | | | | |
| Conditional | Canopy cover | 0.219 | 0.117 | 1.876 | 0.061 | -0.010, 0.447 |
| Conditional | Coarse woody debris | -0.177 | 0.114 | -1.560 | 0.119 | -0.400, 0.045 |
| Zero inflation | Intercept | -2.503 | 0.558 | -4.488 | **<0.001** | -3.596, -1.410 |
| Zero inflation | Site: B vs A | 1.258 | 0.489 | 2.574 | **0.010** | 0.300, 2.216 |
| Zero inflation | Treatment: S[A] vs S[nA] | -0.148 | 0.639 | -0.232 | 0.816 | -1.401, 1.104 |
| Zero inflation | Treatment: G vs S[nA] | 1.915 | 0.575 | 3.330 | **<0.001** | 0.788, 3.041 |
| Random effects | plot_id:site: (Intercept) | 0.553 | | | | |
| Random effects | site: (Intercept) | 0.000 | | | | |
| **Host abundance (across-treatment moisture) — nbinom2 (log link); n = 192; logLik = -480.061; AIC = 990.123** | | | | | | |
| Conditional | Intercept | 1.877 | 0.155 | 12.104 | **<0.001** | 1.573, 2.181 |
| Conditional | Treatment: S[A] vs S[nA] | -0.027 | 0.174 | -0.158 | 0.875 | -0.368, 0.314 |
| Conditional | Treatment: G vs S[nA] | -0.916 | 0.286 | -3.209 | **0.001** | -1.476, -0.357 |
| Conditional | Tree richness | -0.290 | 0.108 | -2.687 | **0.007** | -0.501, -0.078 |
| Conditional | Moisture content (across treatments) | -0.384 | 0.131 | -2.940 | **0.003** | -0.641, -0.128 |
| Conditional | Ant presence | 0.193 | 0.268 | 0.721 | 0.471 | -0.332, 0.718 |
| Conditional | Canopy cover | 0.226 | 0.116 | 1.945 | 0.052 | -0.002, 0.454 |
| Conditional | Coarse woody debris | -0.178 | 0.113 | -1.568 | 0.117 | -0.400, 0.044 |
| Zero inflation | Intercept | -2.485 | 0.554 | -4.484 | **<0.001** | -3.571, -1.399 |
| Zero inflation | Site: B vs A | 1.238 | 0.488 | 2.534 | **0.011** | 0.280, 2.195 |
| Zero inflation | Treatment: S[A] vs S[nA] | -0.127 | 0.630 | -0.202 | 0.840 | -1.363, 1.108 |
| Zero inflation | Treatment: G vs S[nA] | 1.885 | 0.577 | 3.270 | **0.001** | 0.755, 3.016 |
| Random effects | plot_id:site: (Intercept) | 0.552 | | | | |
| Random effects | site: (Intercept) | 0.000 | | | | |
| **Host abundance (design model) — nbinom2 (log link); n = 192; logLik = -482.579; AIC = 987.158** | | | | | | |
| Conditional | Intercept | 1.957 | 0.349 | 5.615 | **<0.001** | 1.274, 2.641 |
| Conditional | Treatment: S[A] vs S[nA] | 0.060 | 0.159 | 0.374 | 0.708 | -0.253, 0.372 |
| Conditional | Treatment: G vs S[nA] | -1.098 | 0.242 | -4.540 | **<0.001** | -1.572, -0.624 |
| Conditional | Tree richness | -0.212 | 0.102 | -2.069 | **0.039** | -0.413, -0.011 |
| Zero inflation | Intercept | -2.460 | 0.538 | -4.572 | **<0.001** | -3.515, -1.405 |
| Zero inflation | Site: B vs A | 1.176 | 0.466 | 2.522 | **0.012** | 0.262, 2.090 |
| Zero inflation | Treatment: S[A] vs S[nA] | -0.063 | 0.617 | -0.102 | 0.919 | -1.272, 1.147 |
| Zero inflation | Treatment: G vs S[nA] | 2.006 | 0.560 | 3.580 | **<0.001** | 0.908, 3.104 |
| Random effects | plot_id:site: (Intercept) | 0.592 | | | | |
| Random effects | site: (Intercept) | 0.448 | | | | |
| **Host richness — genpois (log link); n = 192; logLik = -193.818; AIC = 407.636** | | | | | | |
| Conditional | Intercept | -0.168 | 0.100 | -1.692 | 0.091 | -0.363, 0.027 |
| Conditional | Treatment: S[A] vs S[nA] | 0.040 | 0.098 | 0.411 | 0.681 | -0.151, 0.231 |
| Conditional | Treatment: G vs S[nA] | 0.056 | 0.137 | 0.412 | 0.680 | -0.212, 0.325 |
| Conditional | Moisture content (within treatments) | 0.029 | 0.055 | 0.527 | 0.598 | -0.078, 0.136 |
| Conditional | Tree richness | 0.118 | 0.055 | 2.146 | **0.032** | 0.010, 0.226 |
| Conditional | Canopy cover | -0.078 | 0.057 | -1.383 | 0.167 | -0.190, 0.033 |
| Conditional | Host abundance | 0.861 | 0.076 | 11.274 | **<0.001** | 0.711, 1.011 |
| Conditional | Coarse woody debris | 0.049 | 0.056 | 0.879 | 0.380 | -0.060, 0.158 |
| Random effects | plot_id: (Intercept) | 0.224 | | | | |
| **Host richness (without host abundance) — poisson (log link); n = 192; logLik = -259.012; AIC = 538.024** | | | | | | |
| Conditional | Intercept | 0.370 | 0.108 | 3.424 | **<0.001** | 0.158, 0.582 |
| Conditional | Treatment: S[A] vs S[nA] | -0.017 | 0.144 | -0.121 | 0.904 | -0.299, 0.265 |
| Conditional | Treatment: G vs S[nA] | -1.037 | 0.196 | -5.285 | **<0.001** | -1.422, -0.653 |
| Conditional | Moisture content (within treatments) | -0.266 | 0.078 | -3.419 | **<0.001** | -0.418, -0.113 |
| Conditional | Ant presence | 0.365 | 0.192 | 1.899 | 0.058 | -0.012, 0.742 |
| Conditional | Tree richness | -0.040 | 0.074 | -0.541 | 0.589 | -0.186, 0.106 |
| Conditional | Canopy cover | -0.027 | 0.075 | -0.365 | 0.715 | -0.175, 0.120 |
| Conditional | Coarse woody debris | -0.024 | 0.071 | -0.334 | 0.739 | -0.163, 0.116 |
| Random effects | plot_id:site: (Intercept) | 0.177 | | | | |
| Random effects | site: (Intercept) | 0.000 | | | | |

| Component | Predictor | Estimate | SE | Test statistic | P | 95% CI |
|---|---|---|---|---|---|---|
| **Host richness (design model) — genpois (log link); n = 192; logLik = -257.209; AIC = 536.419** | | | | | | |
| Conditional | Intercept | 0.533 | 0.106 | 5.007 | **<0.001** | 0.324, 0.742 |
| Conditional | Treatment: S[A] vs S[nA] | 0.059 | 0.124 | 0.480 | 0.632 | -0.183, 0.302 |
| Conditional | Treatment: G vs S[nA] | -0.394 | 0.198 | -1.995 | **0.046** | -0.781, -0.007 |
| Conditional | Tree richness | -0.060 | 0.060 | -0.997 | 0.319 | -0.178, 0.058 |
| Zero inflation | Intercept | -3.430 | 0.887 | -3.868 | **<0.001** | -5.167, -1.692 |
| Zero inflation | Site: B vs A | 1.985 | 0.717 | 2.768 | **0.006** | 0.579, 3.390 |
| Zero inflation | Treatment: S[A] vs S[nA] | 0.367 | 0.758 | 0.484 | 0.629 | -1.119, 1.852 |
| Zero inflation | Treatment: G vs S[nA] | 2.379 | 0.735 | 3.236 | **0.001** | 0.938, 3.819 |
| Random effects | plot_id:site: (Intercept) | 0.198 | | | | |
| Random effects | site: (Intercept) | 0.022 | | | | |
| **Parasitoid abundance — nbinom2 (log link); n = 127; logLik = -154.976; AIC = 333.951** | | | | | | |
| Conditional | Intercept | -0.827 | 0.278 | -2.979 | **0.003** | -1.371, -0.283 |
| Conditional | Treatment: S[A] vs S[nA] | 0.133 | 0.283 | 0.469 | 0.639 | -0.422, 0.688 |
| Conditional | Treatment: G vs S[nA] | -0.823 | 0.500 | -1.646 | 0.100 | -1.804, 0.157 |
| Conditional | Moisture content (within treatments) | 0.096 | 0.137 | 0.700 | 0.484 | -0.173, 0.366 |
| Conditional | Ant presence | 0.269 | 0.392 | 0.687 | 0.492 | -0.499, 1.038 |
| Conditional | Tree richness | -0.125 | 0.152 | -0.823 | 0.411 | -0.422, 0.172 |
| Conditional | Canopy cover | 0.451 | 0.160 | 2.809 | **0.005** | 0.136, 0.765 |
| Conditional | Host abundance | 0.824 | 0.213 | 3.869 | **<0.001** | 0.407, 1.241 |
| Conditional | Coarse woody debris | -0.370 | 0.167 | -2.219 | **0.026** | -0.696, -0.043 |
| Random effects | plot_id:site: (Intercept) | 0.000 | | | | |
| Random effects | site: (Intercept) | 0.000 | | | | |
| **Parasitoid abundance (without host abundance) — nbinom2 (log link); n = 127; logLik = -162.407; AIC = 346.815** | | | | | | |
| Conditional | Intercept | -0.128 | 0.221 | -0.581 | 0.562 | -0.561, 0.305 |
| Conditional | Treatment: S[A] vs S[nA] | 0.112 | 0.306 | 0.364 | 0.716 | -0.489, 0.712 |
| Conditional | Treatment: G vs S[nA] | -1.383 | 0.512 | -2.700 | **0.007** | -2.387, -0.379 |
| Conditional | Ant presence | 0.234 | 0.435 | 0.537 | 0.591 | -0.619, 1.087 |
| Conditional | Moisture content (within treatments) | -0.065 | 0.141 | -0.461 | 0.645 | -0.341, 0.211 |
| Conditional | Tree richness | -0.353 | 0.152 | -2.319 | **0.020** | -0.652, -0.055 |
| Conditional | Canopy cover | 0.543 | 0.164 | 3.307 | **<0.001** | 0.221, 0.865 |
| Conditional | Coarse woody debris | -0.368 | 0.172 | -2.141 | **0.032** | -0.705, -0.031 |
| Random effects | plot_id:site: (Intercept) | 0.000 | | | | |
| Random effects | site: (Intercept) | 0.000 | | | | |
| **Parasitoid abundance (design model) — nbinom1 (log link); n = 127; logLik = -169.539; AIC = 353.077** | | | | | | |
| Conditional | Intercept | -0.085 | 0.241 | -0.353 | 0.724 | -0.556, 0.386 |
| Conditional | Treatment: S[A] vs S[nA] | 0.288 | 0.272 | 1.060 | 0.289 | -0.245, 0.820 |
| Conditional | Treatment: G vs S[nA] | -0.962 | 0.494 | -1.948 | 0.051 | -1.929, 0.006 |
| Conditional | Tree richness | -0.134 | 0.142 | -0.946 | 0.344 | -0.413, 0.144 |
| Random effects | plot_id:site: (Intercept) | 0.399 | | | | |
| Random effects | site: (Intercept) | 0.000 | | | | |
| **Parasitoid richness — poisson (log link); n = 127; logLik = -80.171; AIC = 178.343** | | | | | | |
| Conditional | Intercept | -1.503 | 0.271 | -5.538 | **<0.001** | -2.035, -0.971 |
| Conditional | Treatment: S[A] vs S[nA] | 0.109 | 0.273 | 0.398 | 0.691 | -0.427, 0.645 |
| Conditional | Treatment: G vs S[nA] | -0.189 | 0.536 | -0.353 | 0.724 | -1.239, 0.861 |
| Conditional | Ant presence | 0.107 | 0.388 | 0.277 | 0.782 | -0.652, 0.867 |
| Conditional | Moisture content (within treatments) | 0.095 | 0.123 | 0.775 | 0.438 | -0.145, 0.335 |
| Conditional | Tree richness | 0.094 | 0.148 | 0.633 | 0.527 | -0.197, 0.385 |
| Conditional | Canopy cover | -0.030 | 0.151 | -0.200 | 0.841 | -0.326, 0.266 |
| Conditional | Parasitoid abundance | 0.910 | 0.119 | 7.649 | **<0.001** | 0.677, 1.143 |
| Random effects | plot_id: (Intercept) | 0.000 | | | | |
| **Parasitoid richness (without parasitoid abundance) — poisson (log link); n = 127; logLik = -111.988; AIC = 243.976** | | | | | | |
| Conditional | Intercept | -1.064 | 0.254 | -4.188 | **<0.001** | -1.562, -0.566 |
| Conditional | Treatment: S[A] vs S[nA] | 0.247 | 0.258 | 0.958 | 0.338 | -0.258, 0.752 |
| Conditional | Treatment: G vs S[nA] | -0.675 | 0.521 | -1.296 | 0.195 | -1.697, 0.346 |
| Conditional | Host abundance | 0.505 | 0.179 | 2.822 | **0.005** | 0.154, 0.855 |
| Conditional | Ant presence | 0.150 | 0.367 | 0.408 | 0.683 | -0.570, 0.869 |
| Conditional | Moisture content (within treatments) | 0.161 | 0.129 | 1.245 | 0.213 | -0.092, 0.415 |

| Component | Predictor | Estimate | SE | Test statistic | P | 95% CI |
|---|---|---|---|---|---|---|
| **Parasitoid richness (without parasitoid abundance) — poisson (log link); n = 127; logLik = -111.988; AIC = 243.976 (continued)** | | | | | | |
| Conditional | Tree richness | -0.108 | 0.130 | -0.828 | 0.408 | -0.363, 0.147 |
| Conditional | Canopy cover | 0.310 | 0.141 | 2.191 | **0.028** | 0.033, 0.587 |
| Random effects | plot_id:site: (Intercept) | 0.000 | | | | |
| Random effects | site: (Intercept) | 0.000 | | | | |
| **Parasitoid richness (design model) — poisson (log link); n = 127; logLik = -119.308; AIC = 248.616** | | | | | | |
| Conditional | Intercept | -0.601 | 0.189 | -3.177 | **0.001** | -0.972, -0.230 |
| Conditional | Treatment: S[A] vs S[nA] | 0.258 | 0.250 | 1.031 | 0.302 | -0.233, 0.749 |
| Conditional | Treatment: G vs S[nA] | -0.969 | 0.486 | -1.997 | **0.046** | -1.921, -0.018 |
| Conditional | Tree richness | -0.123 | 0.125 | -0.981 | 0.327 | -0.368, 0.123 |
| Random effects | plot_id: (Intercept) | 0.000 | | | | |
| **Apparent parasitism — betabinomial (logit link); n = 127; logLik = -116.521; AIC = 257.041** | | | | | | |
| Conditional | Intercept | -2.713 | 0.342 | -7.942 | **<0.001** | -3.382, -2.043 |
| Conditional | Treatment: S[A] vs S[nA] | 0.020 | 0.299 | 0.068 | 0.945 | -0.565, 0.606 |
| Conditional | Treatment: G vs S[nA] | 0.133 | 0.535 | 0.249 | 0.803 | -0.915, 1.181 |
| Conditional | Moisture content (within treatments) | 0.068 | 0.167 | 0.405 | 0.685 | -0.260, 0.396 |
| Conditional | Ant presence | 0.444 | 0.448 | 0.991 | 0.322 | -0.434, 1.323 |
| Conditional | Parasitoid richness | 1.079 | 0.161 | 6.720 | **<0.001** | 0.764, 1.393 |
| Conditional | Host richness | -0.539 | 0.196 | -2.747 | **0.006** | -0.923, -0.154 |
| Conditional | Tree richness | 0.259 | 0.171 | 1.511 | 0.131 | -0.077, 0.595 |
| Conditional | Canopy cover | -0.133 | 0.194 | -0.686 | 0.492 | -0.514, 0.247 |
| Random effects | plot_id:site: (Intercept) | 0.612 | | | | |
| Random effects | site: (Intercept) | 0.000 | | | | |
| **Apparent parasitism (reduced model) — betabinomial (logit link); n = 127; logLik = -153.617; AIC = 327.234** | | | | | | |
| Conditional | Intercept | -2.194 | 0.255 | -8.593 | **<0.001** | -2.694, -1.693 |
| Conditional | Treatment: S[A] vs S[nA] | 0.275 | 0.304 | 0.906 | 0.365 | -0.321, 0.872 |
| Conditional | Treatment: G vs S[nA] | -0.519 | 0.542 | -0.959 | 0.338 | -1.581, 0.542 |
| Conditional | Moisture content (within treatments) | 0.243 | 0.176 | 1.385 | 0.166 | -0.101, 0.588 |
| Conditional | Ant presence | 0.184 | 0.460 | 0.400 | 0.689 | -0.717, 1.085 |
| Conditional | Tree richness | -0.035 | 0.172 | -0.201 | 0.841 | -0.371, 0.302 |
| Conditional | Canopy cover | 0.359 | 0.193 | 1.865 | 0.062 | -0.018, 0.737 |
| Random effects | plot_id:site: (Intercept) | 0.701 | | | | |
| Random effects | site: (Intercept) | 0.000 | | | | |
| **Apparent parasitism (design model) — betabinomial (logit link); n = 127; logLik = -155.948; AIC = 325.895** | | | | | | |
| Conditional | Intercept | -2.182 | 0.259 | -8.435 | **<0.001** | -2.689, -1.675 |
| Conditional | Treatment: S[A] vs S[nA] | 0.276 | 0.294 | 0.939 | 0.348 | -0.300, 0.852 |
| Conditional | Treatment: G vs S[nA] | -0.493 | 0.524 | -0.940 | 0.347 | -1.520, 0.535 |
| Conditional | Tree richness | 0.065 | 0.174 | 0.374 | 0.708 | -0.277, 0.407 |
| Random effects | plot_id:site: (Intercept) | 0.804 | | | | |
| Random effects | site: (Intercept) | 0.000 | | | | |

*Notes. Estimates are shown on the model link scale. Bold P values are ≤ 0.05. S[nA] is the treatment reference level.*

### C2. PERMANOVA results

Community-composition analyses

| Term | df | Sum of squares | $R^2$ | F | P |
|---|---|---|---|---|---|
| **Deadwood-level host community composition — Bray-Curtis; permutations restricted within plots** | | | | | |
| treatment | 2 | 2.948 | 0.068 | 4.492 | **<0.001** |
| Moisture content (within treatments) | 1 | 0.758 | 0.017 | 2.310 | 0.776 |
| ant_pres | 1 | 0.151 | 0.003 | 0.459 | 0.712 |
| Residual | 120 | 39.385 | 0.906 | | |
| Total | 124 | 43.454 | 1.000 | | |
| **Plot-level host community composition (abundance-based) — Bray-Curtis** | | | | | |
| Tree richness | 1 | 0.548 | 0.029 | 1.750 | 0.070 |
| Canopy cover | 1 | 0.151 | 0.008 | 0.481 | 0.882 |
| Plot-level ant occurrence | 1 | 0.402 | 0.021 | 1.282 | 0.254 |
| Residual | 57 | 17.852 | 0.947 | | |
| Total | 60 | 18.851 | 1.000 | | |
| **Plot-level host community composition (incidence-based) — Binary Bray-Curtis** | | | | | |
| Tree richness | 1 | 0.578 | 0.035 | 2.133 | **0.046** |
| Canopy cover | 1 | 0.164 | 0.010 | 0.605 | 0.733 |
| Plot-level ant occurrence | 1 | 0.330 | 0.020 | 1.217 | 0.312 |
| Residual | 57 | 15.446 | 0.941 | | |
| Total | 60 | 16.413 | 1.000 | | |
| **PERMDISP diagnostic for deadwood treatment — Bray-Curtis** | | | | | |
| Treatment (parametric ANOVA) | 2 | 0.058 | | 0.737 | 0.481 |
| Treatment (permutation test) | 2 | 0.058 | | 0.737 | 0.478 |

*Notes. PERMANOVA tests used 1,000 permutations and marginal sums of squares. PERMDISP reports parametric ANOVA and permutation-test results based on 999 permutations.*

### C3. Structural equation model results

Full structural equation model

| Response | Predictor | Estimate | Std. Estimate | SE | Test statistic | df | P |
|---|---|---|---|---|---|---|---|
| Apparent parasitism | Moisture content | 0.944 | 0.011 | 0.543 | 1.740 | 127 | 0.082 |
| Apparent parasitism | Ant presence | 0.116 | 0.216 | 0.271 | 0.427 | 127 | 0.669 |
| Apparent parasitism | Parasitoid richness | 1.725 | 0.673 | 0.188 | 9.196 | 127 | **<0.001** |
| Apparent parasitism | Host richness | -0.455 | -0.811 | 0.122 | -3.715 | 127 | **<0.001** |
| Apparent parasitism | Tree richness | 0.059 | 0.109 | 0.076 | 0.783 | 127 | 0.433 |
| Apparent parasitism | Canopy cover | -0.694 | -0.039 | 0.382 | -1.818 | 127 | 0.069 |
| Apparent parasitism | Treatment | - | - | - | 4.329 | 2 | 0.115 |
| Apparent parasitism | Treatment: G | -2.225 | 0.427 | 0.269 | -8.261 | Inf | <0.001 |
| Apparent parasitism | Treatment: nA | -2.035 | 0.074 | 0.224 | -9.072 | Inf | <0.001 |
| Apparent parasitism | Treatment: A | -1.705 | 0.037 | 0.187 | -9.122 | Inf | <0.001 |
| Parasitoid richness | Host richness | -0.007 | -0.234 | 0.143 | -0.050 | 118 | 0.960 |
| Parasitoid richness | Host abundance | 0.433 | 0.546 | 0.184 | 2.358 | 118 | **0.018** |
| Parasitoid richness | Ant presence | 0.148 | 0.107 | 0.371 | 0.398 | 118 | 0.691 |
| Parasitoid richness | Moisture content | 0.940 | 0.101 | 0.762 | 1.233 | 118 | 0.218 |
| Parasitoid richness | Tree richness | -0.085 | -0.065 | 0.104 | -0.815 | 118 | 0.415 |
| Parasitoid richness | Canopy cover | 1.081 | 0.194 | 0.514 | 2.102 | 118 | **0.035** |
| Parasitoid richness | Treatment | - | - | - | 5.808 | 2 | 0.055 |
| Parasitoid richness | Treatment: G | -1.795 | -0.434 | 0.471 | -3.808 | Inf | <0.001 |
| Parasitoid richness | Treatment: nA | -0.920 | -0.132 | 0.304 | -3.022 | Inf | 0.003 |
| Parasitoid richness | Treatment: A | -0.665 | 0.087 | 0.261 | -2.547 | Inf | 0.011 |
| Host richness | Moisture content | 0.168 | 0.013 | 0.304 | 0.552 | 192 | 0.581 |
| Host richness | Tree richness | 0.093 | 0.090 | 0.044 | 2.107 | 192 | **0.035** |
| Host richness | Canopy cover | -0.245 | -0.041 | 0.198 | -1.242 | 192 | 0.214 |
| Host richness | Host abundance | 0.732 | 0.896 | 0.065 | 11.346 | 192 | **<0.001** |
| Host richness | Treatment | - | - | - | 0.154 | 2 | 0.926 |
| Host richness | Treatment: nA | -0.155 | 0.009 | 0.100 | -1.552 | Inf | 0.121 |
| Host richness | Treatment: G | -0.139 | -0.004 | 0.123 | -1.127 | Inf | 0.260 |
| Host richness | Treatment: A | -0.117 | -0.004 | 0.099 | -1.179 | Inf | 0.238 |
| Host abundance | Tree richness | -0.077 | -0.089 | 0.032 | -2.439 | 192 | **0.015** |
| Host abundance | Moisture content | -0.712 | -0.300 | 0.227 | -3.141 | 192 | **0.002** |
| Host abundance | Ant presence | 0.063 | 0.163 | 0.109 | 0.575 | 192 | 0.566 |
| Host abundance | Canopy cover | 0.192 | 0.050 | 0.153 | 1.254 | 192 | 0.210 |
| Host abundance | Treatment | - | - | - | 8.360 | 2 | **0.015** |
| Host abundance | Treatment: G | 0.375 | -0.419 | 0.090 | 4.168 | Inf | **<0.001** |
| Host abundance | Treatment: A | 0.653 | 0.301 | 0.072 | 9.078 | Inf | **<0.001** |
| Host abundance | Treatment: nA | 0.683 | 0.292 | 0.079 | 8.698 | Inf | **<0.001** |
| Moisture content | Tree richness | -0.077 | -0.057 | 0.050 | -1.541 | 173 | 0.123 |
| Moisture content | Canopy cover | 0.368 | 0.063 | 0.243 | 1.513 | 173 | 0.130 |
| Moisture content | Treatment | - | - | - | 129.589 | 2 | **<0.001** |
| Moisture content | Treatment: A | -1.107 | -0.353 | 0.510 | -2.168 | Inf | **0.030** |
| Moisture content | Treatment: nA | -1.101 | -0.329 | 0.510 | -2.158 | Inf | **0.031** |
| Moisture content | Treatment: G | 0.002 | 0.687 | 0.516 | 0.004 | Inf | 0.997 |
| Ant presence | Tree richness | 0.277 | 0.348 | 0.203 | 1.367 | 192 | 0.172 |
| Ant presence | Canopy cover | 1.068 | 0.303 | 1.045 | 1.022 | 192 | 0.307 |
| Ant presence | Treatment | - | - | - | 8.266 | 2 | **0.016** |
| Ant presence | Treatment: nA | -3.510 | -3.510 | 0.753 | -4.660 | Inf | **<0.001** |
| Ant presence | Treatment: A | -2.053 | -2.053 | 0.498 | -4.124 | Inf | **<0.001** |
| Ant presence | Treatment: G | -1.464 | -1.464 | 0.420 | -3.483 | Inf | **<0.001** |
| Canopy cover | Tree richness | 0.072 | 0.317 | 0.025 | 2.892 | 64 | **0.004** |
| Ant presence (correlated error) | Moisture content | 0.099 | 0.164 | - | 1.371 | 192 | 0.086 |
| Host abundance (correlated error) | Apparent parasitism | -0.122 | -0.119 | - | -1.689 | 192 | **0.046** |

*Notes. SE, test statistic, df, and P are from the response-specific SEM. Standardization-model estimates are standardized coefficients only for continuous-to-continuous paths. Treatment and ant-presence coefficients retain their contrast or log-odds interpretation.*

### C3. Structural equation model results

Parsimonious structural equation model

| Response | Predictor | Estimate | Std. Estimate | SE | Test statistic | df | P |
|---|---|---|---|---|---|---|---|
| Apparent parasitism | Moisture content | 0.958 | 0.042 | 0.543 | 1.763 | 127 | 0.078 |
| Apparent parasitism | Parasitoid richness | 1.697 | 0.662 | 0.185 | 9.156 | 127 | **<0.001** |
| Apparent parasitism | Host richness | -0.446 | -0.799 | 0.122 | -3.642 | 127 | **<0.001** |
| Apparent parasitism | Canopy cover | -0.557 | 0.002 | 0.346 | -1.608 | 127 | 0.108 |
| Apparent parasitism | Treatment | - | - | - | 4.224 | 2 | 0.121 |
| Apparent parasitism | Treatment: G | -2.242 | 0.349 | 0.267 | -8.401 | Inf | <0.001 |
| Apparent parasitism | Treatment: nA | -2.067 | -0.026 | 0.191 | -10.809 | Inf | <0.001 |
| Apparent parasitism | Treatment: A | -1.736 | -0.031 | 0.163 | -10.676 | Inf | <0.001 |
| Parasitoid richness | Host abundance | 0.465 | 0.452 | 0.144 | 3.222 | 121 | **0.001** |
| Parasitoid richness | Moisture content | 0.961 | 0.101 | 0.761 | 1.262 | 121 | 0.207 |
| Parasitoid richness | Canopy cover | 0.987 | 0.183 | 0.484 | 2.038 | 121 | **0.042** |
| Parasitoid richness | Treatment | - | - | - | 5.564 | 2 | 0.062 |
| Parasitoid richness | Treatment: G | -1.824 | -0.560 | 0.464 | -3.931 | Inf | <0.001 |
| Parasitoid richness | Treatment: nA | -1.016 | -0.279 | 0.246 | -4.126 | Inf | <0.001 |
| Parasitoid richness | Treatment: A | -0.740 | -0.041 | 0.233 | -3.174 | Inf | 0.002 |
| Host richness | Tree richness | 0.092 | 0.090 | 0.044 | 2.087 | 192 | **0.037** |
| Host richness | Canopy cover | -0.245 | -0.043 | 0.197 | -1.243 | 192 | 0.214 |
| Host richness | Host abundance | 0.721 | 0.892 | 0.061 | 11.869 | 192 | **<0.001** |
| Host abundance | Tree richness | -0.074 | -0.075 | 0.031 | -2.389 | 192 | **0.017** |
| Host abundance | Moisture content | -0.708 | -0.249 | 0.226 | -3.135 | 192 | **0.002** |
| Host abundance | Canopy cover | 0.195 | 0.038 | 0.153 | 1.272 | 192 | 0.203 |
| Host abundance | Treatment | - | - | - | 8.003 | 2 | **0.018** |
| Host abundance | Treatment: G | 0.363 | -0.497 | 0.088 | 4.144 | Inf | **<0.001** |
| Host abundance | Treatment: A | 0.634 | 0.263 | 0.064 | 9.930 | Inf | **<0.001** |
| Host abundance | Treatment: nA | 0.655 | 0.234 | 0.062 | 10.625 | Inf | **<0.001** |
| Moisture content | Tree richness | -0.077 | -0.057 | 0.050 | -1.541 | 173 | 0.123 |
| Moisture content | Canopy cover | 0.368 | 0.063 | 0.243 | 1.513 | 173 | 0.130 |
| Moisture content | Treatment | - | - | - | 129.589 | 2 | **<0.001** |
| Moisture content | Treatment: A | -1.107 | -0.353 | 0.510 | -2.168 | Inf | **0.030** |
| Moisture content | Treatment: nA | -1.101 | -0.329 | 0.510 | -2.158 | Inf | **0.031** |
| Moisture content | Treatment: G | 0.002 | 0.687 | 0.516 | 0.004 | Inf | 0.997 |
| Ant presence | Tree richness | 0.277 | 0.348 | 0.203 | 1.367 | 192 | 0.172 |
| Ant presence | Canopy cover | 1.068 | 0.303 | 1.045 | 1.022 | 192 | 0.307 |
| Ant presence | Treatment | - | - | - | 8.266 | 2 | **0.016** |
| Ant presence | Treatment: nA | -3.510 | -3.510 | 0.753 | -4.660 | Inf | **<0.001** |
| Ant presence | Treatment: A | -2.053 | -2.053 | 0.498 | -4.124 | Inf | **<0.001** |
| Ant presence | Treatment: G | -1.464 | -1.464 | 0.420 | -3.483 | Inf | **<0.001** |
| Canopy cover | Tree richness | 0.072 | 0.317 | 0.025 | 2.892 | 64 | **0.004** |
| Ant presence (correlated error) | Moisture content | 0.099 | 0.164 | - | 1.371 | 192 | 0.086 |
| Host abundance (correlated error) | Apparent parasitism | -0.117 | -0.094 | - | -1.616 | 192 | 0.054 |

*Notes. SE, test statistic, df, and P are from the response-specific SEM. Standardization-model estimates are standardized coefficients only for continuous-to-continuous paths. Treatment and ant-presence coefficients retain their contrast or log-odds interpretation.*
