## Appendix D for "Moisture limits the diversity of nesting bees, wasps, and parasitoids in lying and standing deadwood"

### Appendix D. Sensitivity model results

Sensitivity model results for the manuscript: "Moisture limits the diversity of nesting bees, wasps, and parasitoids in lying and standing deadwood"

Massimo Martini, Matteo Dadda, Felix Fornoff, Heike Feldhaar, Arong Luo, Finn Rehling, Joshua E. Spitz, Michael Staab, Simon Thorn, Chao-Dong Zhu, Alexandra-Maria Klein

D1. Host-diversity sensitivity models (p. 1)

D2. Alternative ant-variable sensitivity models (p. 2)

D3. Complete-case sensitivity models (p. 3-4)

#### D1. Host-diversity sensitivity models

Model coefficient tables

| Component | Predictor | Estimate | SE | Test statistic | P | 95% CI |
|---|---|---|---|---|---|---|
| **Host richness standardized to 90% coverage — gaussian (identity link); n = 123; logLik = -9.526; AIC = 39.052** | | | | | | |
| Conditional | Intercept | 0.847 | 0.044 | 19.290 | **<0.001** | 0.761, 0.933 |
| Conditional | Treatment: S[A] vs S[nA] | 0.017 | 0.053 | 0.315 | 0.753 | -0.088, 0.121 |
| Conditional | Treatment: G vs S[nA] | 0.021 | 0.073 | 0.285 | 0.776 | -0.122, 0.163 |
| Conditional | Moisture content (within treatments) | 0.026 | 0.027 | 0.970 | 0.332 | -0.027, 0.080 |
| Conditional | Ant presence | 0.082 | 0.070 | 1.168 | 0.243 | -0.056, 0.220 |
| Conditional | Tree richness | 0.028 | 0.025 | 1.132 | 0.257 | -0.020, 0.077 |
| Conditional | Canopy cover | -0.041 | 0.026 | -1.573 | 0.116 | -0.092, 0.010 |
| Conditional | Host abundance | 0.186 | 0.036 | 5.127 | **<0.001** | 0.115, 0.257 |
| Random effects | plot_id: (Intercept) | 0.000 | | | | |
| Random effects | Residual: Observation | 0.261 | | | | |
| **Host Shannon diversity — tweedie (log link); n = 192; logLik = -104.333; AIC = 230.665** | | | | | | |
| Conditional | Intercept | -1.874 | 0.233 | -8.059 | **<0.001** | -2.329, -1.418 |
| Conditional | Treatment: S[A] vs S[nA] | 0.060 | 0.246 | 0.244 | 0.807 | -0.422, 0.542 |
| Conditional | Treatment: G vs S[nA] | 0.236 | 0.369 | 0.640 | 0.522 | -0.487, 0.960 |
| Conditional | Moisture content (within treatments) | 0.092 | 0.130 | 0.707 | 0.479 | -0.163, 0.347 |
| Conditional | Tree richness | 0.184 | 0.112 | 1.642 | 0.101 | -0.036, 0.403 |
| Conditional | Canopy cover | -0.135 | 0.119 | -1.133 | 0.257 | -0.368, 0.098 |
| Conditional | Host abundance | 1.069 | 0.152 | 7.044 | **<0.001** | 0.772, 1.367 |
| Conditional | Coarse woody debris | 0.122 | 0.121 | 1.001 | 0.317 | -0.116, 0.360 |
| Random effects | plot_id: (Intercept) | 0.000 | | | | |
| **Host Shannon diversity (without host abundance) — tweedie (log link); n = 192; logLik = -130.579; AIC = 285.158** | | | | | | |
| Conditional | Intercept | -1.153 | 0.184 | -6.260 | **<0.001** | -1.513, -0.792 |
| Conditional | Treatment: S[A] vs S[nA] | 0.035 | 0.256 | 0.137 | 0.891 | -0.467, 0.538 |
| Conditional | Treatment: G vs S[nA] | -1.018 | 0.339 | -2.999 | **0.003** | -1.683, -0.353 |
| Conditional | Moisture content (within treatments) | -0.248 | 0.128 | -1.935 | 0.053 | -0.499, 0.003 |
| Conditional | Ant presence | 0.462 | 0.322 | 1.434 | 0.151 | -0.169, 1.094 |
| Conditional | Tree richness | -0.027 | 0.123 | -0.220 | 0.826 | -0.269, 0.214 |
| Conditional | Canopy cover | -0.057 | 0.124 | -0.456 | 0.648 | -0.299, 0.186 |
| Conditional | Coarse woody debris | 0.006 | 0.117 | 0.047 | 0.962 | -0.223, 0.235 |
| Random effects | plot_id:site: (Intercept) | 0.000 | | | | |
| Random effects | site: (Intercept) | 0.000 | | | | |

*Notes. Estimates are shown on the model link scale. Bold P values are ≤ 0.05. S[nA] is the treatment reference level.*

### D2. Alternative ant-variable sensitivity models

Model coefficient tables

| Component | Predictor | Estimate | SE | Test statistic | P | 95% CI |
|---|---|---|---|---|---|---|
| **Host occurrence: deadwood-level ant abundance — binomial (logit link); n = 192; logLik = -97.713; AIC = 207.425** | | | | | | |
| Conditional | Intercept | 2.231 | 0.456 | 4.889 | **<0.001** | 1.337, 3.125 |
| Conditional | Treatment: S[A] vs S[nA] | 0.101 | 0.464 | 0.217 | 0.828 | -0.809, 1.011 |
| Conditional | Treatment: G vs S[nA] | -2.109 | 0.473 | -4.456 | **<0.001** | -3.036, -1.181 |
| Conditional | Site: B vs A | -1.406 | 0.395 | -3.557 | **<0.001** | -2.181, -0.631 |
| Conditional | Deadwood-level ant abundance | 0.038 | 0.189 | 0.203 | 0.839 | -0.332, 0.408 |
| Random effects | plot_id: (Intercept) | 0.198 | | | | |
| **Host occurrence: plot-level ant activity — binomial (logit link); n = 192; logLik = -97.226; AIC = 206.452** | | | | | | |
| Conditional | Intercept | 2.219 | 0.456 | 4.866 | **<0.001** | 1.325, 3.112 |
| Conditional | Treatment: S[A] vs S[nA] | 0.107 | 0.464 | 0.232 | 0.817 | -0.802, 1.017 |
| Conditional | Treatment: G vs S[nA] | -2.111 | 0.473 | -4.461 | **<0.001** | -3.039, -1.184 |
| Conditional | Site: B vs A | -1.376 | 0.393 | -3.496 | **<0.001** | -2.147, -0.604 |
| Conditional | Plot-level ant occurrence | 0.186 | 0.187 | 0.994 | 0.320 | -0.180, 0.552 |
| Random effects | plot_id: (Intercept) | 0.165 | | | | |
| **Host abundance: deadwood-level ant abundance — nbinom2 (log link); n = 192; logLik = -480.002; AIC = 990.005** | | | | | | |
| Conditional | Intercept | 2.008 | 0.151 | 13.290 | **<0.001** | 1.712, 2.304 |
| Conditional | Treatment: S[A] vs S[nA] | 0.010 | 0.171 | 0.058 | 0.954 | -0.325, 0.345 |
| Conditional | Treatment: G vs S[nA] | -1.221 | 0.260 | -4.692 | **<0.001** | -1.731, -0.711 |
| Conditional | Tree richness | -0.276 | 0.108 | -2.550 | **0.011** | -0.489, -0.064 |
| Conditional | Moisture content (within treatments) | -0.330 | 0.109 | -3.030 | **0.002** | -0.543, -0.116 |
| Conditional | Deadwood-level ant abundance | 0.015 | 0.094 | 0.164 | 0.869 | -0.169, 0.200 |
| Conditional | Canopy cover | 0.220 | 0.117 | 1.888 | 0.059 | -0.008, 0.448 |
| Conditional | Coarse woody debris | -0.176 | 0.113 | -1.555 | 0.120 | -0.399, 0.046 |
| Zero inflation | Intercept | -2.504 | 0.558 | -4.490 | **<0.001** | -3.597, -1.411 |
| Zero inflation | Site: B vs A | 1.252 | 0.487 | 2.573 | **0.010** | 0.298, 2.205 |
| Zero inflation | Treatment: S[A] vs S[nA] | -0.133 | 0.637 | -0.209 | 0.834 | -1.383, 1.116 |
| Zero inflation | Treatment: G vs S[nA] | 1.937 | 0.574 | 3.374 | **<0.001** | 0.812, 3.062 |
| Random effects | plot_id:site: (Intercept) | 0.551 | | | | |
| Random effects | site: (Intercept) | 0.000 | | | | |
| **Host abundance: plot-level ant activity — nbinom2 (log link); n = 192; logLik = -478.559; AIC = 987.117** | | | | | | |
| Conditional | Intercept | 2.005 | 0.145 | 13.806 | **<0.001** | 1.720, 2.289 |
| Conditional | Treatment: S[A] vs S[nA] | 0.018 | 0.167 | 0.110 | 0.912 | -0.309, 0.346 |
| Conditional | Treatment: G vs S[nA] | -1.183 | 0.253 | -4.683 | **<0.001** | -1.678, -0.688 |
| Conditional | Tree richness | -0.293 | 0.104 | -2.820 | **0.005** | -0.496, -0.089 |
| Conditional | Moisture content (within treatments) | -0.287 | 0.108 | -2.655 | **0.008** | -0.499, -0.075 |
| Conditional | Plot-level ant occurrence | 0.182 | 0.105 | 1.743 | 0.081 | -0.023, 0.387 |
| Conditional | Canopy cover | 0.200 | 0.113 | 1.767 | 0.077 | -0.022, 0.421 |
| Conditional | Coarse woody debris | -0.221 | 0.114 | -1.947 | 0.051 | -0.444, 0.001 |
| Zero inflation | Intercept | -2.529 | 0.562 | -4.496 | **<0.001** | -3.631, -1.426 |
| Zero inflation | Site: B vs A | 1.276 | 0.484 | 2.637 | **0.008** | 0.328, 2.224 |
| Zero inflation | Treatment: S[A] vs S[nA] | -0.095 | 0.631 | -0.150 | 0.881 | -1.332, 1.142 |
| Zero inflation | Treatment: G vs S[nA] | 1.974 | 0.574 | 3.440 | **<0.001** | 0.849, 3.098 |
| Random effects | plot_id:site: (Intercept) | 0.515 | | | | |
| Random effects | site: (Intercept) | 0.000 | | | | |

*Notes. Estimates are shown on the model link scale. Bold P values are ≤ 0.05. S[nA] is the treatment reference level.*

### D3. Complete-case sensitivity models

Model coefficient tables

| Component | Predictor | Estimate | SE | Test statistic | P | 95% CI |
|---|---|---|---|---|---|---|
| **Host abundance: complete cases — nbinom2 (log link); n = 173; logLik = -448.499; AIC = 926.997** | | | | | | |
| Conditional | Intercept | 2.019 | 0.155 | 13.036 | **<0.001** | 1.715, 2.322 |
| Conditional | Treatment: S[A] vs S[nA] | -0.033 | 0.180 | -0.185 | 0.853 | -0.386, 0.319 |
| Conditional | Treatment: G vs S[nA] | -1.285 | 0.286 | -4.500 | **<0.001** | -1.844, -0.725 |
| Conditional | Tree richness | -0.271 | 0.114 | -2.382 | **0.017** | -0.494, -0.048 |
| Conditional | Moisture content (within treatments) | -0.366 | 0.118 | -3.095 | **0.002** | -0.598, -0.134 |
| Conditional | Ant presence | 0.198 | 0.273 | 0.724 | 0.469 | -0.338, 0.734 |
| Conditional | Canopy cover | 0.170 | 0.124 | 1.376 | 0.169 | -0.072, 0.413 |
| Conditional | Coarse woody debris | -0.158 | 0.121 | -1.302 | 0.193 | -0.395, 0.080 |
| Zero inflation | Intercept | -2.778 | 0.703 | -3.954 | **<0.001** | -4.155, -1.401 |
| Zero inflation | Site: B vs A | 1.042 | 0.567 | 1.836 | 0.066 | -0.070, 2.153 |
| Zero inflation | Treatment: S[A] vs S[nA] | -0.009 | 0.842 | -0.011 | 0.991 | -1.660, 1.641 |
| Zero inflation | Treatment: G vs S[nA] | 2.280 | 0.717 | 3.181 | **0.001** | 0.875, 3.685 |
| Random effects | plot_id:site: (Intercept) | 0.585 | | | | |
| Random effects | site: (Intercept) | 0.000 | | | | |
| **Host richness: complete cases — genpois (log link); n = 173; logLik = -179.473; AIC = 378.946** | | | | | | |
| Conditional | Intercept | -0.166 | 0.105 | -1.577 | 0.115 | -0.373, 0.040 |
| Conditional | Treatment: S[A] vs S[nA] | 0.063 | 0.097 | 0.650 | 0.516 | -0.128, 0.254 |
| Conditional | Treatment: G vs S[nA] | 0.071 | 0.139 | 0.509 | 0.611 | -0.202, 0.344 |
| Conditional | Moisture content (within treatments) | 0.022 | 0.057 | 0.393 | 0.694 | -0.089, 0.134 |
| Conditional | Tree richness | 0.107 | 0.057 | 1.886 | 0.059 | -0.004, 0.219 |
| Conditional | Canopy cover | -0.082 | 0.061 | -1.363 | 0.173 | -0.201, 0.036 |
| Conditional | Host abundance | 0.833 | 0.078 | 10.653 | **<0.001** | 0.679, 0.986 |
| Conditional | Coarse woody debris | 0.040 | 0.059 | 0.675 | 0.500 | -0.076, 0.155 |
| Random effects | plot_id: (Intercept) | 0.249 | | | | |
| **Host richness (without host abundance): complete cases — poisson (log link); n = 173; logLik = -235.847; AIC = 491.693** | | | | | | |
| Conditional | Intercept | 0.423 | 0.111 | 3.801 | **<0.001** | 0.205, 0.641 |
| Conditional | Treatment: S[A] vs S[nA] | -0.031 | 0.148 | -0.211 | 0.833 | -0.321, 0.258 |
| Conditional | Treatment: G vs S[nA] | -1.035 | 0.204 | -5.070 | **<0.001** | -1.435, -0.635 |
| Conditional | Moisture content (within treatments) | -0.268 | 0.080 | -3.351 | **<0.001** | -0.425, -0.111 |
| Conditional | Ant presence | 0.378 | 0.191 | 1.980 | **0.048** | 0.004, 0.753 |
| Conditional | Tree richness | -0.049 | 0.075 | -0.655 | 0.512 | -0.195, 0.097 |
| Conditional | Canopy cover | -0.064 | 0.075 | -0.849 | 0.396 | -0.212, 0.084 |
| Conditional | Coarse woody debris | -0.019 | 0.072 | -0.260 | 0.795 | -0.160, 0.122 |
| Random effects | plot_id:site: (Intercept) | 0.139 | | | | |
| Random effects | site: (Intercept) | 0.000 | | | | |
| **Parasitoid abundance: complete cases — nbinom2 (log link); n = 120; logLik = -149.660; AIC = 323.320** | | | | | | |
| Conditional | Intercept | -0.830 | 0.293 | -2.836 | **0.005** | -1.404, -0.256 |
| Conditional | Treatment: S[A] vs S[nA] | 0.160 | 0.297 | 0.540 | 0.589 | -0.421, 0.742 |
| Conditional | Treatment: G vs S[nA] | -0.731 | 0.517 | -1.415 | 0.157 | -1.744, 0.281 |
| Conditional | Moisture content (within treatments) | 0.097 | 0.147 | 0.662 | 0.508 | -0.190, 0.384 |
| Conditional | Ant presence | 0.237 | 0.402 | 0.588 | 0.556 | -0.551, 1.024 |
| Conditional | Tree richness | -0.108 | 0.157 | -0.690 | 0.490 | -0.415, 0.199 |
| Conditional | Canopy cover | 0.436 | 0.167 | 2.617 | **0.009** | 0.109, 0.763 |
| Conditional | Host abundance | 0.825 | 0.221 | 3.738 | **<0.001** | 0.392, 1.257 |
| Conditional | Coarse woody debris | -0.362 | 0.173 | -2.090 | **0.037** | -0.701, -0.023 |
| Random effects | plot_id:site: (Intercept) | 0.000 | | | | |
| Random effects | site: (Intercept) | 0.000 | | | | |
| **Parasitoid abundance (without host abundance): complete cases — nbinom2 (log link); n = 120; logLik = -156.582; AIC = 335.165** | | | | | | |
| Conditional | Intercept | -0.109 | 0.233 | -0.469 | 0.639 | -0.566, 0.347 |
| Conditional | Treatment: S[A] vs S[nA] | 0.130 | 0.321 | 0.404 | 0.686 | -0.500, 0.759 |
| Conditional | Treatment: G vs S[nA] | -1.278 | 0.531 | -2.409 | **0.016** | -2.318, -0.238 |
| Conditional | Ant presence | 0.197 | 0.446 | 0.442 | 0.659 | -0.677, 1.070 |
| Conditional | Moisture content (within treatments) | -0.067 | 0.150 | -0.449 | 0.654 | -0.361, 0.227 |

| Component | Predictor | Estimate | SE | Test statistic | P | 95% CI |
|---|---|---|---|---|---|---|
| **Parasitoid abundance (without host abundance): complete cases — nbinom2 (log link); n = 120; logLik = -156.582; AIC = 335.165 (continued)** | | | | | | |
| Conditional | Tree richness | -0.336 | 0.157 | -2.138 | **0.033** | -0.643, -0.028 |
| Conditional | Canopy cover | 0.518 | 0.170 | 3.041 | **0.002** | 0.184, 0.852 |
| Conditional | Coarse woody debris | -0.350 | 0.179 | -1.957 | **0.050** | -0.700, 0.001 |
| Random effects | plot_id:site: (Intercept) | 0.000 | | | | |
| Random effects | site: (Intercept) | 0.000 | | | | |
| **Parasitoid richness: complete cases — poisson (log link); n = 120; logLik = -76.029; AIC = 170.058** | | | | | | |
| Conditional | Intercept | -1.548 | 0.287 | -5.391 | **<0.001** | -2.111, -0.985 |
| Conditional | Treatment: S[A] vs S[nA] | 0.135 | 0.281 | 0.479 | 0.632 | -0.416, 0.686 |
| Conditional | Treatment: G vs S[nA] | -0.104 | 0.541 | -0.192 | 0.848 | -1.163, 0.956 |
| Conditional | Ant presence | 0.084 | 0.389 | 0.216 | 0.829 | -0.679, 0.846 |
| Conditional | Moisture content (within treatments) | 0.096 | 0.128 | 0.751 | 0.453 | -0.155, 0.348 |
| Conditional | Tree richness | 0.116 | 0.150 | 0.778 | 0.437 | -0.177, 0.410 |
| Conditional | Canopy cover | -0.029 | 0.154 | -0.191 | 0.849 | -0.332, 0.273 |
| Conditional | Parasitoid abundance | 0.918 | 0.122 | 7.526 | **<0.001** | 0.679, 1.156 |
| Random effects | plot_id: (Intercept) | 0.000 | | | | |
| **Parasitoid richness (without parasitoid abundance): complete cases — poisson (log link); n = 120; logLik = -107.730; AIC = 235.459** | | | | | | |
| Conditional | Intercept | -1.081 | 0.264 | -4.099 | **<0.001** | -1.597, -0.564 |
| Conditional | Treatment: S[A] vs S[nA] | 0.267 | 0.264 | 1.012 | 0.311 | -0.250, 0.785 |
| Conditional | Treatment: G vs S[nA] | -0.575 | 0.527 | -1.092 | 0.275 | -1.608, 0.457 |
| Conditional | Host abundance | 0.500 | 0.181 | 2.761 | **0.006** | 0.145, 0.854 |
| Conditional | Ant presence | 0.122 | 0.369 | 0.329 | 0.742 | -0.602, 0.845 |
| Conditional | Moisture content (within treatments) | 0.164 | 0.135 | 1.215 | 0.224 | -0.101, 0.429 |
| Conditional | Tree richness | -0.096 | 0.130 | -0.736 | 0.462 | -0.351, 0.159 |
| Conditional | Canopy cover | 0.305 | 0.145 | 2.109 | **0.035** | 0.022, 0.588 |
| Random effects | plot_id:site: (Intercept) | 0.000 | | | | |
| Random effects | site: (Intercept) | 0.000 | | | | |
| **Apparent parasitism: complete cases — betabinomial (logit link); n = 120; logLik = -111.463; AIC = 246.926** | | | | | | |
| Conditional | Intercept | -2.760 | 0.357 | -7.734 | **<0.001** | -3.459, -2.060 |
| Conditional | Treatment: S[A] vs S[nA] | 0.080 | 0.312 | 0.255 | 0.798 | -0.531, 0.691 |
| Conditional | Treatment: G vs S[nA] | 0.248 | 0.548 | 0.452 | 0.651 | -0.826, 1.321 |
| Conditional | Moisture content (within treatments) | 0.040 | 0.179 | 0.224 | 0.823 | -0.311, 0.392 |
| Conditional | Ant presence | 0.416 | 0.457 | 0.912 | 0.362 | -0.479, 1.312 |
| Conditional | Parasitoid richness | 1.100 | 0.167 | 6.585 | **<0.001** | 0.773, 1.428 |
| Conditional | Host richness | -0.572 | 0.202 | -2.830 | **0.005** | -0.969, -0.176 |
| Conditional | Tree richness | 0.285 | 0.176 | 1.621 | 0.105 | -0.060, 0.630 |
| Conditional | Canopy cover | -0.168 | 0.202 | -0.832 | 0.405 | -0.563, 0.227 |
| Random effects | plot_id:site: (Intercept) | 0.620 | | | | |
| Random effects | site: (Intercept) | 0.000 | | | | |
| **Apparent parasitism (reduced model): complete cases — betabinomial (logit link); n = 120; logLik = -147.838; AIC = 315.676** | | | | | | |
| Conditional | Intercept | -2.220 | 0.268 | -8.274 | **<0.001** | -2.746, -1.694 |
| Conditional | Treatment: S[A] vs S[nA] | 0.320 | 0.316 | 1.013 | 0.311 | -0.299, 0.940 |
| Conditional | Treatment: G vs S[nA] | -0.391 | 0.555 | -0.704 | 0.482 | -1.478, 0.697 |
| Conditional | Moisture content (within treatments) | 0.255 | 0.189 | 1.349 | 0.177 | -0.116, 0.626 |
| Conditional | Ant presence | 0.130 | 0.470 | 0.277 | 0.782 | -0.790, 1.050 |
| Conditional | Tree richness | -0.022 | 0.178 | -0.125 | 0.900 | -0.371, 0.326 |
| Conditional | Canopy cover | 0.346 | 0.201 | 1.721 | 0.085 | -0.048, 0.739 |
| Random effects | plot_id:site: (Intercept) | 0.730 | | | | |
| Random effects | site: (Intercept) | 0.000 | | | | |

*Notes. Estimates are shown on the model link scale. Bold P values are ≤ 0.05. S[nA] is the treatment reference level.*
